# Time-Resolved Proteomics of a Polyglutamine-Expanded Huntingtin in vitro Cell Model Reveals Early RNA-Regulatory Compensation and Late Respiratory-Chain Remodeling

**DOI:** 10.64898/2026.08.21.746221

**Authors:** Ekin Sonmez, Pinar Mutlu, Cinla Ozlevent, Mehmet Sarihan, Gurler Akpinar, Murat Kasap, Huseyin Cimen

**Affiliations:** The Institute of Biotechnology, Gebze Technical University, Kocaeli, Turkiye; Department of Molecular Biology and Genetics, Gebze Technical University, Kocaeli, Turkiye; Department of Medical Biology, Faculty of Medicine, Kocaeli University, Kocaeli, Turkiye; Central Research Laboratory Application and Research Center (GTUMAR), Gebze Technical University, Kocaeli, Turkiye

**Keywords:** Huntington’s disease, polyglutamine, quantitative proteomics, temporal dynamics, mitochondrial dysfunction, oxidative phosphorylation, proteostasis, drug repurposing

## Abstract

Huntington’s disease (HD) is caused by a polyglutamine-expanded huntingtin protein that exerts progressive cellular toxicity. However, the temporal order of pathogenic events, and in particular which changes arise early and which emerge late, remains incompletely defined, despite their distinct therapeutic implications. To examine this trajectory, we profiled the proteome of a huntingtin-expressing cell model at early (72 h) and late (144 h) stages. Rather than a linear progression, the proteome changed in two distinct stages. At the early stage, cells showed increased abundance of RNA-processing, splicing, and protein-synthesis proteins, which may reflect an adaptive response to preserve gene-expression fidelity under stress. By the late stage, many of these proteins were depleted and mitochondrial energy-metabolism proteins dominated the response. Notably, 85% of proteins altered at both time points reversed direction between stages (95% at a false-discovery rate below 5%), and all reversals were supported by a significant genotype-by-time interaction, indicating that mutant huntingtin reorganizes the proteome rather than amplifying a fixed set of perturbations. Subunit-level analysis of the respiratory chain showed that Complex IV and ATP synthase subunits were reduced at 72 h, whereas at 144 h subunits of all five complexes were broadly increased, most strongly in Complex I and III, with a few distal subunits (COX5B, ATP5F1A, ATP5F1B) remaining reduced. Leveraging this proteomic map, we applied an AI-assisted, direction-aware drug repurposing strategy. Of 1,712 differentially abundant proteins, 498 were druggable, 109 had at least one agent whose mechanism was concordant with the required correction, and 89 of these 109 proteins mapped to at least one agent listed as approved in the source databases (56 to an agent with current regulatory approval). These included the approved Complex I inhibitor metformin, the investigational Complex I inhibitor ME-344, and pathway-level mitochondria-directed agents (elamipretide/SS-31, approved only for Barth syndrome; MitoQ, a dietary supplement), several of which have previously been evaluated in HD models. Collectively, these findings describe a two-stage proteomic response to mutant huntingtin in this cell model and suggest a putative early window for preclinical intervention testing, before late respiratory-chain remodeling is established.

## Introduction

Huntington’s disease (HD) is an autosomal-dominant neurodegenerative disorder caused by an expanded CAG trinucleotide repeat in the first exon of the HTT gene, which is translated into an elongated polyglutamine (polyQ) tract within the huntingtin protein (Huntington’s Disease Collaborative Research Group, 1993). Expansions beyond a threshold of approximately 35-40 glutamines confer conformational instability and a propensity to adopt β-sheet-rich, amyloid-like structures, driving the formation of intracellular inclusions that are a pathological hallmark of the disease (Scherzinger et al., 1997; DiFiglia et al., 1997; Ross and Poirier, 2004). Although these inclusions were long considered the primary cytotoxic entity, prefibrillar oligomeric species are now increasingly recognized as the more proximal drivers of toxicity, capable of perturbing proteostasis, mitochondrial function and intracellular transport before terminal inclusion bodies form (Arrasate et al., 2004; Ross and Tabrizi, 2011; Bates et al., 2015; Saudou and Humbert, 2016).

Mutant huntingtin exerts pleiotropic effects on cellular physiology as documented among patient tissue and experimental models like transcriptional dysregulation, impaired RNA metabolism, disruption of the nuclear pore complex, mitochondrial dysfunction and progressive failure of protein-quality-control networks have each been (Zuccato et al., 2010; Cui et al., 2006; Labadorf et al., 2015; Grima et al., 2017). In particular, mitochondrial bioenergetic failure including transcriptional repression of PGC-1α and a measurable cellular energy deficit is a recurrent theme in HD pathophysiology (Cui et al., 2006; Mochel and Haller, 2011), and recent syntheses place mitochondrial and proteostatic collapse at the center of molecular pathogenesis (Tabrizi et al., 2020). However, most studies capture a single disease stage, and the temporal order in which these processes emerge is rarely resolved. Distinguishing early, potentially adaptive responses from later changes is important, because the two may imply different therapeutic strategies. Global proteomics is well suited to reconstructing this temporal architecture, as it measures protein abundance, the molecular layer closest to cellular function, and captures compensatory as well as degenerative programs. A recent proteomic profiling of HD medium spiny neurons has unraveled the value of this approach (Tshilenge et al., 2023). While neurodegenerative diseases are fundamentally characterized by neuronal vulnerability, the initial molecular events triggering proteostasis imbalance are often obscured in highly specialized cellular contexts. In this study, the HEK293T cell model was used to study the cell-autonomous kinetics of polyglutamine (polyQ) aggregation and the associated proteotoxic stress, separately from neuron-specific secondary effects. This non-neuronal system allows for a highly controlled, reproducible evaluation of the temporal relationship between mutant huntingtin accumulation, early RNA-regulatory compensation, and late mitochondrial changes, free from the confounding variables of synaptic or glial interactions (Carnemolla et al., 2017; Yablonska et al., 2019). HEK293T cells also express several neuronal-lineage proteins, including neurofilament subunits (Shaw et al., 2002). Our aim was to define the proteomic and metabolic changes that accompany the formation of polyQ aggregates in a controlled cellular system.

So, we use a transient HEK293T model of polyQ-expanded huntingtin to build a time-resolved proteomic map of mutant-driven proteome remodeling. By profiling the mutant (Htt-Q74) and wild-type (Htt-Q23) proteomes at an early (72 h) and a late (144 h) time point and applying a temporal-classification framework, we follow the proteome trajectory.

HD drug development has repeatedly targeted energy metabolism through antioxidant, mitochondrial and bioenergetic agents yet major trials have been largely negative (Huntington Study Group, 2001; Hersch et al., 2017). We propose a time-resolved, direction-aware proteomic map to resolve this compromise. We therefore coupled our map to a systematic, AI-assisted target-nomination process using a large language model research agent (Claude Science, beta, San Francisco, CA, USA, underlying model Claude Opus 4.5) (Anthropic, 2026) to intersect the differentially abundant proteins with established resources and to retain only agents whose mechanism of action would correct rather than reinforce the observed direction of change. This direction-aware framework distinguishes our approach from prior energy-focused repurposing attempts and yields a stage-matched shortlist of candidates that can be tested within the putative early window suggested by the proteomic trajectory.

## 2 Materials and Methods

### 2.1 Cell culture and transfection

HEK293T cells were cultured in high-glucose DMEM (Gibco, USA) supplemented with 10% fetal bovine serum (FBS; Gibco) and 1% penicillin/streptomycin (Gibco) at 37°C and 5% CO₂. For transfection, the medium was replaced with transfection mixture containing plasmid DNA (12 µg) and PEIpro reagent (1:1) per 1.5 × 10⁶ cells. Cells were transfected with a GFP control, wild-type Htt-Q23, or mutant Htt-Q74 construct (GFP-tagged, Addgene #114492) and documented by using fluorescence microscopy (BioRad, ZOE Fluorescent Cell Imager) (Yiğit et al., 2023). The GFP-only control was used for model validation (fluorescence microscopy and immunoblotting) and was not included in the proteomic analysis; all proteomic comparisons were Htt-Q74 (MT) versus Htt-Q23 (WT) at the same time point.

For aggregation analysis, GFP fluorescence of Htt-Q23 and Htt-Q74 cells was imaged at 72 h and 144 h post-transfection on an inverted fluorescence microscope (Leica DMi8; 10× and 40× objectives). Images were analysed in ImageJ/Fiji: the GFP signal was segmented by u Dark thresholding after background subtraction (rolling ball radius: 50 px; particle size filter 2 px²– infinity), and the area and circularity (4π × area / perimeter²) of each segmented object were measured. Cell nuclei were counterstained with Hoechst 33342, pre-processed (rolling ball radius = 30 px, contrast enhancement saturated at 0.35, and smooth filtering), and segmented using Huang Dark thresholding followed by a watershed binary separation with a size filter of 20 px²–infinity. Aggregate load per cell was calculated as the number of puncta divided by the number of cells and expressed relative to 72 h. Values are mean ± SD of n ≥ 18 fields per group from 3 independent transfections; Q23 and Q74 were compared with a two-tailed unpaired Student’s t-test, as were 72 h and 144 h.

### 2.2 Immunoblotting

Cell lysates prepared in a RIPA buffer with protease/phosphatase inhibitors (1×) were clarified (14,000 *g*, 15 min, 4°C) and protein concentration was determined by BCA assay. Proteins from cell lysates (30 µg) were denatured in Laemmli buffer with β-mercaptoethanol (95°C, 5 min) and separated with SDS-PAGE. Protein samples were then transferred to methanol-activated PVDF membranes (Trans-Blot, Bio-Rad). This was followed with membrane blocking in skim milk/TBS-T (5% for 1 h at room temperature), and probing overnight at 4°C with anti-GFP primary antibody (1:15000 in 5% BSA, E-AB-48010, Elabscience). Membranes were incubated with corresponding HRP-conjugated secondary antibodies (1:5000, 1 h, at room temperature) and their images were recorded on a iBright750 system (Invitrogen). Chemiluminescent signals were quantified in ImageJ (Gel Analyzer) and normalized to β-actin (Üstüner and Çimen, 2016; Arı Uyar et al., 2024; Yiğit et al., 2026).

### 2.3 Mass spectrometry-based proteomic analyses

Protein samples were processed using the filter-aided sample preparation (FASP) method (Wiśniewski et al., 2009; Rencber et al., 2024). For each experimental condition, proteins were extracted from five independent cell cultures, and equal protein amounts were combined to generate a single pooled sample. Each pooled sample was analysed in three technical LC-MS/MS runs to assess analytical reproducibility. The pooling strategy was employed to obtain a representative composite protein sample for each experimental condition. However, because only one biological pool was prepared per condition, variability among the individual cultures could not be estimated, and the three runs per group are technical rather than independent biological replicates. The DEqMS differential-abundance analysis (Section 2.5) was therefore used as an exploratory screen to select candidate proteins: its p-values reflect analytical rather than biological variability and are not interpreted as evidence of statistically validated biological differences, and the log₂ fold changes describe differences between pooled samples. The candidate proteins require confirmation in independent biological samples.

For digestion, 200 µg of protein was mixed with 200 µL of 8 M urea in 100 mM Tris and applied to Microcon Ultracel 30-kDa filter units (Millipore). After tryptic digestion, peptides were eluted in 50 mM ammonium bicarbonate/0.5 M NaCl (14,000 *g*), dried (SpeedVac, Eppendorf), and reconstituted in 0.1% formic acid (FA). Peptide concentrations were determined by Qubit assay (Invitrogen, Q33211) and analyzed on an Ultimate 3000 RSLC nano system (Dionex/Thermo) coupled to a Q-Exactive mass spectrometer (Thermo). Mobile phases were 0.1% FA (A) and 80% acetonitrile/0.1% FA (B). Peptides were pre-concentrated and desalted on a trap column and separated over a 120 min multi-step gradient (6-90% B) at 300 nL/min. Data-acquisition parameters followed Sarihan et al. (Sarihan et al., 2024).

### 2.4 Database search and data analysis

Raw spectra were converted to open mzML format (ProteoWizard MSConvert, peak-picking filter, vendor centroiding) and searched against the human UniProt reference proteome (canonical + isoforms, plus a contaminant panel and reversed decoys) in FragPipe (v24.0) using the MSFragger (v4.4.1) search engine (Figure 1) (Kong et al., 2017). Precursor and fragment mass tolerances were set to 20 ppm with mass calibration and parameter optimization enabled. Trypsin was specified with up to three missed cleavages, peptide length was restricted to 6-50 residues and peptide mass to 500-5000 Da. Deep-learning-based rescoring was performed with MSBooster (predicting retention time, spectra, and ion mobility) (Yang et al., 2023), and peptide-spectrum matches were re-scored and validated with Percolator (v3.7.1) (Käll et al., 2007). Protein inference and statistical filtering were performed in Philosopher (v5.1.3) (da Veiga Leprevost et al., 2020), with peptide-and protein-level identifications filtered to 1% false-discovery rate (FDR). Only proteins supported by at least two unique peptides were retained. Label-free quantification used IonQuant (v1.11.20) with the MaxLFQ algorithm and match-between-runs (MBR) enabled (Yu et al., 2021). Each analytical run was assigned to its experiment group (wild-type or mutant) as required for MBR, and unique+razor peptides were used for protein-level roll-up.

**Figure 1.**
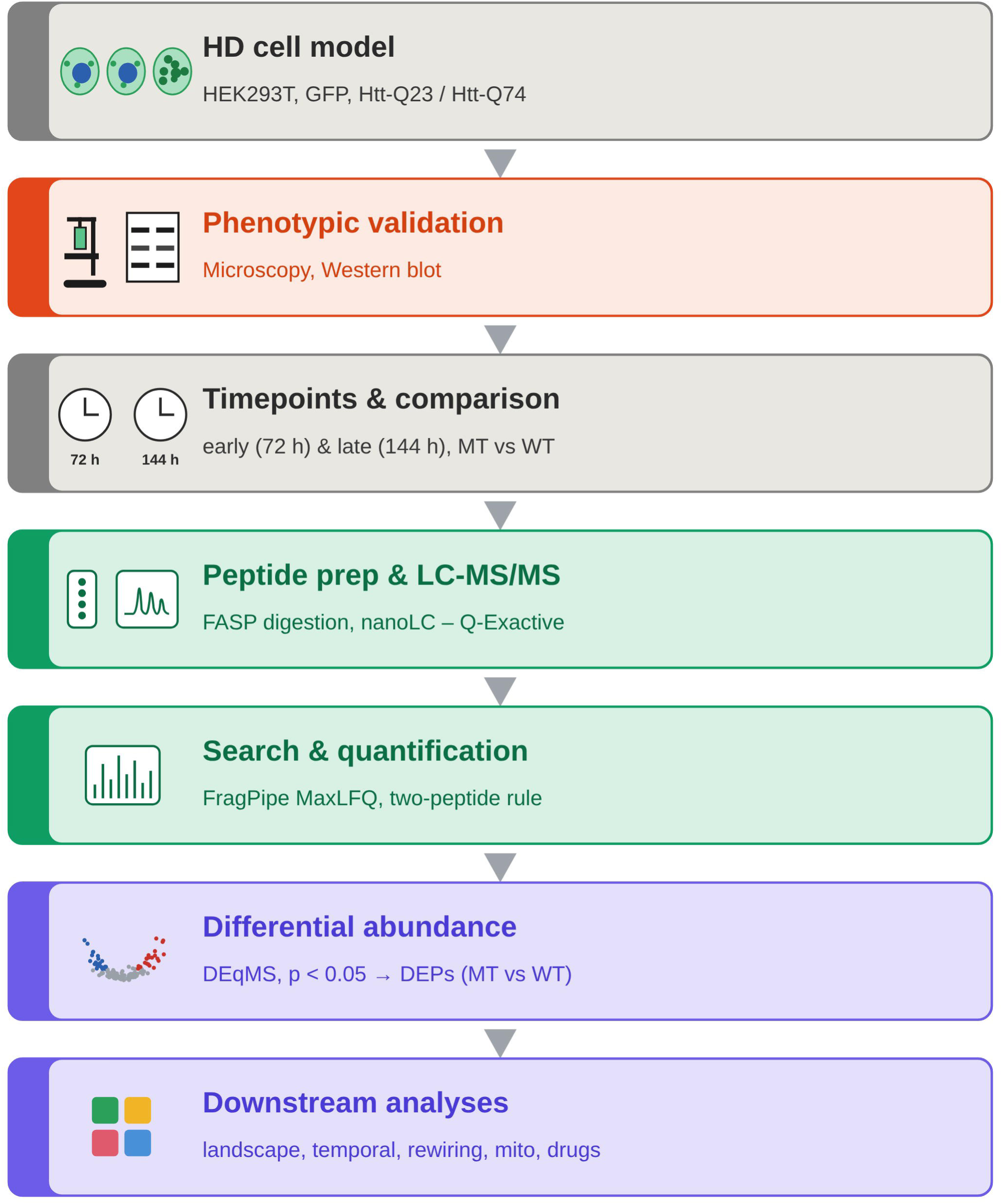
Experimental and analytical workflow of the temporal huntingtin proteomics study. Schematic overview of the study design. HEK293T cells were transiently transfected with a GFP control, wild-type Htt-Q23 or mutant Htt-Q74 construct and validated phenotypically by fluorescence microscopy and western blotting (GFP control used for validation only). Two time points capturing the early (72 h) and late (144 h) response were selected for label-free quantitative proteomics. Peptides were prepared by FASP, analysed by nanoLC-MS/MS (Q-Exactive), searched and quantified in FragPipe (MSFragger/IonQuant, MaxLFQ, two-peptide rule) and tested for differential abundance (MT vs WT) with DEqMS (p < 0.05). Downstream analyses comprised the global proteomic landscape, temporal architecture, functional rewiring, respiratory-chain remodeling and direction-aware drug repurposing.

### 2.5 Label-free quantification and differential expression

Protein-level MaxLFQ intensities were extracted from the FragPipe combined_protein output, log₂-transformed, and normalization methods were compared with NormalyzerDE. Median-centred normalization was selected on the basis of intensity-distribution, coefficient-of-variation, and PCA quality metrics (Willforss et al., 2019). Missing values were handled by a hybrid strategy that distinguishes missingness mechanisms. For proteins absent from an entire condition (missing-not-at-random, MNAR), missing values were drawn for each protein from a normal distribution centred on the 1st percentile of that protein’s observed intensities, with the protein’s observed standard deviation, whereas sporadic missing values (missing-at-random, MAR) were imputed by k-nearest-neighbours (k = 5). Proteins undetected across all samples were removed (Supplementary Table S1). Differential abundance between wild-type (WT, Htt-Q23) and mutant (MT, Htt-Q74) was evaluated separately at each time point using DEqMS, which augments the limma empirical-Bayes framework with a peptide/spectral-count-dependent variance model to avoid overconfident calls for low-evidence proteins (Zhu et al., 2020); the combined total spectral count of each protein across the six runs of the corresponding time point (FragPipe) was used as the count variable. Results were robust to using the peptide count instead (Supplementary Table S2). Proteins with an unadjusted DEqMS p < 0.05 were defined as differentially abundant proteins (DEPs); no fold-change cut-off was applied. Benjamini–Hochberg-adjusted p-values (FDR; Benjamini and Hochberg, 1995) are reported for all proteins (Supplementary Tables S1 and S2). A sensitivity analysis excluding proteins with imputed values was also performed (Supplementary Table S2). Direction of change was calculated from the MT/WT log₂ fold-change. Separate MT-versus-WT contrasts were tested at 72 h and at 144 h (Supplementary Table S2).

To test directly whether the genotype effect changed over time, a joint linear model with genotype, time and a genotype-by-time interaction term was fitted with limma and DEqMS, using the same spectral-count-based variance moderation as the main analysis (Ritchie et al., 2015; Zhu et al., 2020), for proteins quantified at both time points (n = 2,266), and interaction p-values were adjusted by the Benjamini–Hochberg method.

### 2.6 Temporal classification and functional analysis

DEPs were intersected across time points and classified as early-specific (72h only), late-specific (144h only) or persistent/shared (Supplementary Table S2). For shared DEPs, directional stability was scored as progressive (same sign, |144h| > |72h |), stable ( same sign, |144h| ≤ |72h|) or reversed (opposite sign between time points). Reversal was evaluated only for proteins detected as DEPs at both time points. Over-representation analysis (GO:BP and KEGG) was performed with clusterProfiler and org.Hs.eg.db (Wu et al., 2021), using the genome-wide annotation as background and Benjamini–Hochberg correction (adjusted p < 0.05). As a sensitivity analysis, enrichment was repeated with FDR-filtered DEP lists and with all proteins quantified at either time point as the background (Supplementary Figure 6; Supplementary Table S3). Term membership was drawn from database annotations rather than from the enrichment results themselves.

For the pathway-trajectory analysis (Figure 3D; Supplementary Figure 3), GO:BP and KEGG over-representation was run separately for the early-specific, shared and late-specific DEP sets (gene-set size 5–800), and terms with an adjusted p < 0.05 in at least one set and assigned to metabolism or proteostasis by keyword were retained. For each term and set, the directionality score at each time point was the mean log₂FC (MT/WT) of the term’s member genes that were DEPs at that time point, and the Δ score was the 144 h score minus the 72 h score; scores were then averaged across the sets in which the term was enriched. The Functional Rewiring Index of a functional category was the mean |Δ score| of its terms (Supplementary Table S4). For the drug-repurposing analysis, reversed proteins were further labelled as ‘collapse’ (increased at 72 h and decreased at 144 h) or ‘rebound’ (decreased at 72 h and increased at 144 h).

Mitochondrial proteins and their sub-organellar localization, OXPHOS complex membership and redox annotation were assigned using MitoCarta 3.0 (Rath et al., 2021). All primary figures were derived from DEqMS-filtered DEP lists (p < 0.05). Raw-intensity analyses were used only as orthogonal validation. UniProt accessions served as the join key, with UNIPROT and SYMBOL conversion via ClusterProfiler bitr and a UniProt REST fallback.

### 2.7 Drug-repurposing analysis

Candidate therapeutic targets were nominated from the differential-expression tables with a systematic, AI-assisted, direction-aware pipeline executed with an autonomous large language model research agent (Claude Science, beta, San Francisco, CA, USA, underlying model Claude Opus 4.5, Anthropic 2026) (Anthropic, 2026) (Figure 1). For every DEP, the agent queried established resources (DGIdb, Open Targets, DrugBank and ChEMBL) and recorded, for each gene, the supporting resources, an integer-evidence count (0-4), candidate drugs (prioritizing approved and clinical-stage agents, highest development phase first), their mechanism of action, and the highest clinical phase reached. Drug names, mechanisms and identifiers were taken only from these verified database records. The agent was explicitly constrained not to invent drug-target links, and genes without database support were retained but marked non-druggable. Each gene’s trajectory class and directions were supplied from the temporal-classification step and used as given. A direction-aware corrective logic was then applied and for proteins elevated in mutant cells the corrective action was defined as SUPPRESS and for depleted proteins as RESTORE. Each candidate agent was labelled mechanism-concordant only when its pharmacology (e.g. inhibitor/antagonist versus activator/agonist) matched the required correction. Candidates were organized into two layers (Figure 5B). Layer 1 was target-anchored agents acting on a specific druggable DEP and Layer 2 was pathway-level mitochondrial agents (e.g. SS-31/elamipretide, MitoQ, idebenone) that act on bioenergetic or redox processes rather than on a single protein target and they were included on the basis of curated HD/energy-metabolism literature. Candidates were tiered by trajectory class and evidence strength (Tier 1-3) (Supplementary Table S6). Clinical phase was recorded at the target level as the highest phase reached by any candidate agent for that target; the regulatory status of individual agents named in the text (approved, investigational, preclinical or dietary supplement) was verified separately against regulatory sources and ClinicalTrials.gov (November 2026); for mechanism-concordant targets, the current approval status of each candidate agent is reported in Supplementary Table S6.

## 3 Results

### 3.1 Temporal aggregation kinetics of polyQ-expanded huntingtin

HEK293T cells transfected with GFP and Htt-Q23 (WT) maintained a diffuse cytoplasmic and nuclear signal at both time points. In contrast, cells transfected with Htt-Q74 (MT) contained discrete intracellular puncta characteristic of mutant-huntingtin aggregates at both time points (Supplementary Figure 1A, B). Quantitative image analysis showed that the GFP signal in Q23 cells covered large, irregular areas corresponding to whole cell bodies (segmented signal area ∼160–200 px², circularity ∼0.65–0.70), whereas Q74 cells contained small, round puncta (∼55– 80 px², circularity ∼0.85; p < 0.001 for both parameters at 72 h and 144 h; Supplementary Figure 1D). The number of puncta per cell was approximately halved at 144 h relative to 72 h, while cell number approximately doubled and the total integrated GFP signal remained unchanged (Supplementary Figure 1C); whether this reflects loss of aggregate-bearing cells or dilution by proliferating cells cannot be distinguished with the present data. The two proteomic time points therefore represent an early stage at which aggregates have formed (72 h) and a later stage with a lower per-cell aggregate load in an expanding cell population (144 h). Expression of the GFP-tagged constructs at the two time points selected for proteomic profiling, 72 h (early) and 144 h (late), was confirmed by anti-GFP immunoblotting (Supplementary Figure 1E, F).

### 3.2 Global proteomic landscape reveals a marked early-to-late shift

Label-free quantification of nanoLC-MS/MS data followed by principal component analysis separated samples primarily by genotype along PC1 (43.1%) and by time along PC2 (27.1%), with samples clustering by group and a mutation-associated divergence that intensified over time (Figure 2A). Differential analysis (MT vs WT; DEqMS p < 0.05) identified 412 up-and 379 down-regulated proteins at 72 h (2,579 quantified proteins, Figure 2C) and a substantially larger response at 144 h, with 699 up- and 741 down-regulated proteins (2,500 quantified, Figure 2D). At FDR < 0.05, 356 proteins (208 up- and 148 down-regulated) and 1,266 proteins (617 up- and 649 down-regulated) remained significant at 72 h and 144 h, respectively (Supplementary Tables S1 and S2). The log₂-fold-change distribution broadened and became more left-skewed at 144 h relative to 72 h (Figure 2B), consistent with a shift toward global protein depletion. Among the strongest early changes were up-regulation of RNA-processing and translation factors (SRRM2, EIF4H, EIF4B, UBAP2L, BCLAF1, SERBP1) and down-regulation of quality-control and chromatin-associated proteins (HECTD1, NEMF, ZNF638, FUS, TOP2A) (Figure 2C). SRRM2 is a core scaffolding component of nuclear speckles that organizes co-transcriptional splicing (Ilik et al., 2020), EIF4H and EIF4B are translation-initiation factors whose activity gates aberrant protein synthesis in repeat-expansion disease (Goodman et al., 2019), and both FUS and the ribosome-associated quality-control factor NEMF are directly tied to the RNA binding protein, stress granule and co-translational surveillance pathways whose disruption is a recurrent axis of neurodegeneration (Patel et al., 2015; Shao et al., 2015; Filbeck et al., 2022; Gutiérrez-Garcia et al., 2023). By 144 h, many of these same RNA regulatory proteins (EIF4H, SERBP1, UBAP2L, BCLAF1, HNRNPK, MATR3) were instead depleted, while nuclear and chromatin-remodeling factors (SSRP1, SUPT16H, NUP210) and the mitochondrial phosphate carrier (Solute carrier family 25 member 3, SLC25A3) were elevated (Figure 2D). Matrin 3 (MATR3) and related prion-like-domain RNA-binding proteins are established contributors to proteotoxic RNP granule pathology (Picchiarelli and Dupuis, 2020). Over-representation analysis (GO:BP and KEGG) across the up- and down-regulated sets at each time point showed that RNA splicing, biogenesis of ribonucleoprotein complexes, cytoplasmic translation, and nucleocytoplasmic transport dominated the early response (Supplementary Figure 2A, B). At the late stage, respiratory-chain and oxidative phosphorylation terms became prominent among increased proteins (Supplementary Figure 2A, B). Together, these enrichment patterns indicate that an early RNA-regulatory and translational signature is followed by a late bioenergetic one (Supplementary Table S3).

**Figure 2.**
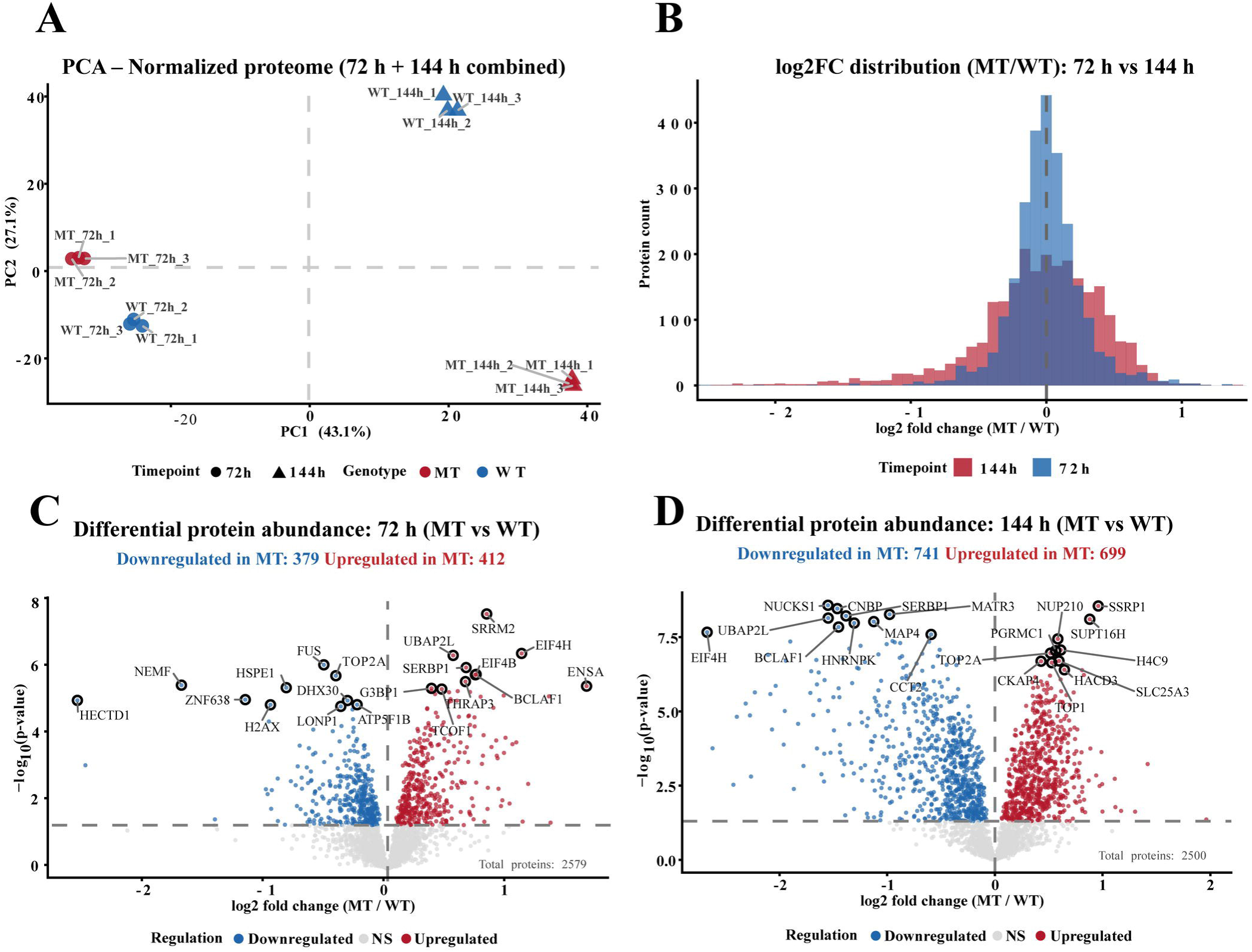
Global proteomic landscape of HEK293T cells expressing mutant (Htt-Q74) versus wild-type (Htt-Q23) huntingtin at 72 h and 144. **h.** (A) Principal component analysis of the normalized proteome (72 h + 144 h combined); PC1 (43.1%) separates genotype and PC2 (27.1%) separates time point. MT, red; WT, blue; circles = 72 h, triangles = 144 h; n = 3 technical replicates per group. (B) Distribution of log₂ fold-change (MT/WT) at 72 h (blue) versus 144 h (red); the 144 h distribution is broader and left-skewed, consistent with a shift toward global protein depletion. (C) Volcano plot of differential protein abundance at 72 h (MT vs WT; DEqMS p < 0.05; 2,579 quantified proteins): 412 proteins up-regulated and 379 down-regulated in MT. (D) Volcano plot at 144 h (MT vs WT; p < 0.05; 2,500 quantified proteins): 699 up-regulated and 741 down-regulated in MT. In (C) and (D) blue = down-regulated, red = up-regulated, grey = not significant; the horizontal dashed line marks p = 0.05 and the top-ranked DEPs by significance are labelled.

### 3.3 The mutant proteome undergoes large-scale directional reversal

To resolve how the proteome shifts for the mutant-expressing cells, DEPs were intersected across time points (Figure 3A). Of all DEPs, only 519 were shared between 72 h and 144 h, whereas 272 were early-specific (72 h only) and 921 were late-specific (144 h only), corresponding to 30%, 16% and 54% of the DEP pool, respectively (Figure 3B). The UpSet decomposition showed that the largest single categories were late-specific up- and down-regulated proteins, with a prominent 72 h-up and 144 h-down intersection indicative of directional reversal (Figure 3A). Analysis of the 519 shared DEPs revealed that the dominant behavior was reversal with 85% (n = 439) changed sign between the two time points; however, only 10% (n = 54) were progressive and 5% (n = 26) were stable (Figure 3B). This pattern was robust to multiple-testing correction: at FDR < 0.05, 222 of 234 shared proteins (95%) reversed direction (Supplementary Table S2). In a joint genotype-by-time model, all 439 reversed proteins showed a significant interaction term (FDR < 0.05) in the expected direction, confirming that their genotype effect changed between time points. The reversal was also robust to excluding imputed values (440 of 520 shared proteins, 85%) and to using peptide instead of spectral counts in DEqMS (438 of 514, 85%) (Supplementary Table S2). This large-scale reversal, which was reported as the anti-diagonal enrichment in the time-progression scatter (Figure 3C), indicates that the mutant proteome is reorganized between time points rather than linearly amplified. Faceted classification of the shared, early-specific and late-specific sets, with the leading genes per subclass labelled (Progressive_up n = 19, Progressive_down n = 35, Stable_up n = 12, Stable_down n = 14; Figure 3C), placed RNA-regulatory and translation-related proteins (EIF4H, RPL35A, NOLC1, YLPM1) among the most strongly reversed consistent with the nucleolar and ribosomal stress that accompanies mutant-huntingtin expression (Lee et al., 2014; Eshraghi et al., 2021) and identified mitochondrial proteins (MRPL3, SLC25A12, UQCRFS1, MGST3) among the progressive changes (Contreras, 2015; Cahill et al., 2020).

**Figure 3.**
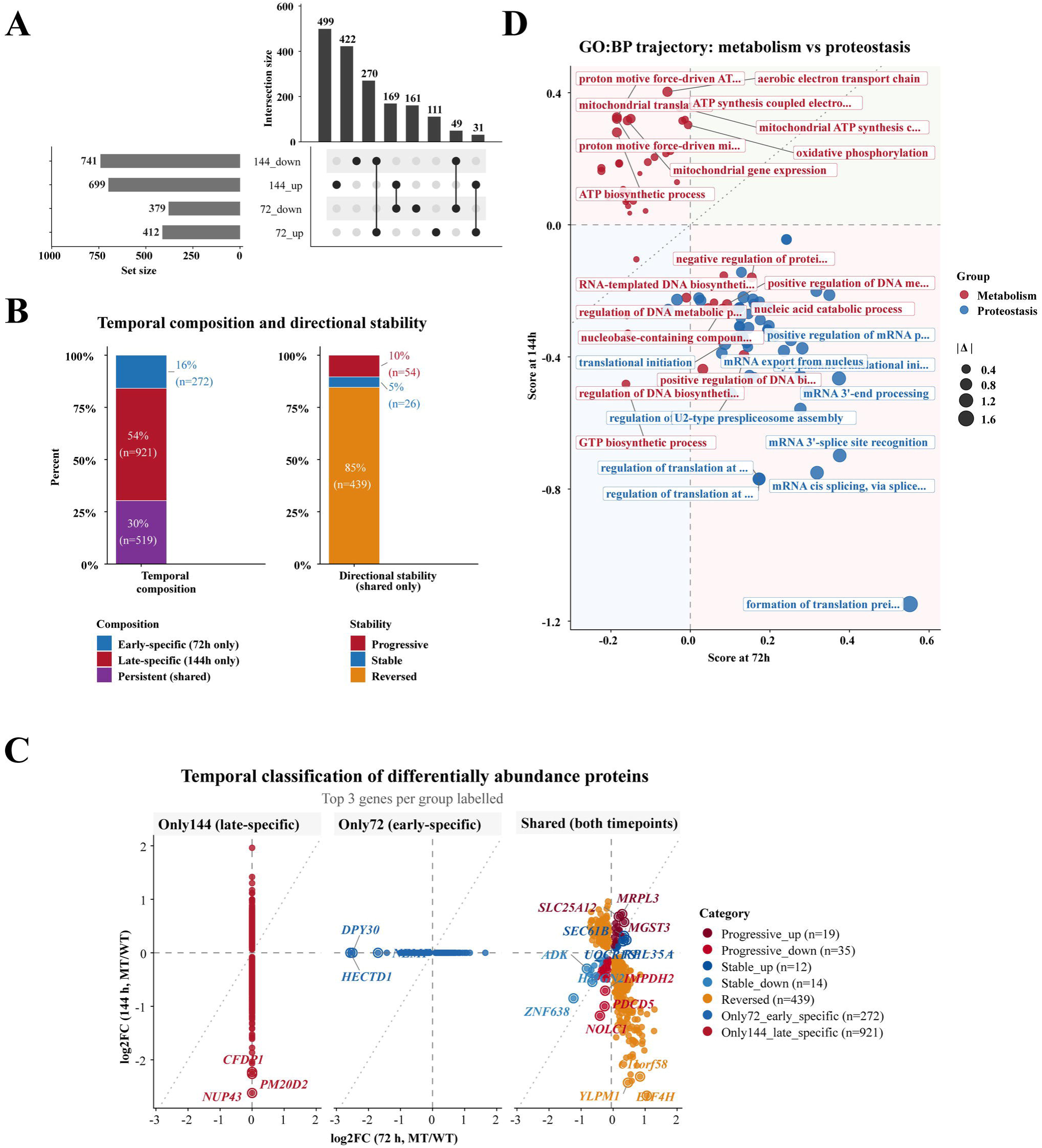
Temporal architecture of the mutant proteome: large-scale directional reversal at the protein and pathway level. (**A**) UpSet plot of DEP overlaps between time points; set sizes are 144_down = 741, 144_up = 699, 72_down = 379 and 72_up = 412, and the intersection bars show that the largest shared category corresponds to proteins that change sign between time points. (**B**) Temporal composition of all DEPs, early-specific (72 h only) 16% (n = 272), late-specific (144 h only) 54% (n = 921) and persistent/shared 30% (n = 519), together with the directional stability of the shared set: 85% reversed (n = 439), 10% progressive (n = 54) and 5% stable (n = 26). (**C**) Temporal classification of DEPs, faceted by class (Only144/late-specific n = 921; Only72/early-specific n = 272; Sftared n = 519) and plotted as log₂FC at 72 ft versus log₂FC at 144 ft. Points are coloured by progression class (Progressive_up n = 19, Progressive_down n = 35, Stable_up n = 12, Stable_down n = 14, Reversed n = 439); single-time-point proteins are plotted on tfte corresponding zero axis and tfte top tftree genes per group are labelled. Tfte dotted diagonals mark conservation (y = x) and reversal (y = −x). (**D**) GO:BP pathway-trajectory plot: each enriched term is positioned by its directionality score at 72 h (x-axis) versus 144 h (y-axis) and coloured by module (metabolism, red; proteostasis, blue); point size scales with |Δ score|. Metabolic and proteostasis terms occupy opposite quadrants, indicating that the two modules move in opposite directions between the early and late stage.

The reversal of directional change was evaluated exclusively for proteins classified as differentially abundant (DEPs) at both time points. Proteins that were dysregulated early and returned to baseline (i.e. no longer significant at 144 h) were captured as early-specific rather than as reciprocal changes. Both behaviors were consistent with the same underlying interpretation that early responses were not maintained and the early-specific and the populations exhibiting reversed trajectories collectively account for the transient nature of the initial response. Functional enrichment of the temporal clusters was in line with this reading: early-specific and reversed proteins were enriched for spliceosome, ribosome biogenesis and nucleocytoplasmic transport, whereas late-specific and progressive changes were enriched for oxidative phosphorylation and related energy-metabolism terms, and the persistent cluster also mapped to the KEGG Huntington’s and Parkinson’s disease pathways (Supplementary Figure 2C, D). In a sensitivity analysis with FDR-filtered lists and the quantified proteome as background, the main terms were retained (RNA splicing, spliceosome and cytoplasmic translation among proteins increased at 72 h and in the shared set, and RNA splicing among proteins decreased at 144 h; oxidative phosphorylation and respiration-related terms among proteins increased at 144 h), whereas the KEGG Huntington’s disease pathway was no longer enriched and early-specific proteins showed no significant terms (Supplementary Figure 6; complete results in Supplementary Table S3).

### 3.4 Functional rewiring dissociates metabolism from proteostasis

To characterize the functional logic of this reprogramming, we compared the temporal trajectories of metabolism and proteostasis associated with GO:BP terms (Figure 3D). In the pathway-trajectory plot, each term is positioned by its enrichment score at 72 h versus 144 h and colored by module (Figure 3D). The two modules occupied opposite quadrants, metabolic processes including proton motive force driven ATP synthesis, mitochondrial translation, and mitochondrial gene expression showed decreased member-protein abundance at 72 h and increased abundance by 144 h. Whereas proteostasis-related processes, cytoplasmic translational pre-initiation complex formation and initiation, mRNA 3’-splice-site recognition, U2-type prespliceosome assembly, mRNA cis-splicing, and mRNA export from the nucleus showed increased member-protein abundance at the early stage and decreased abundance at the later stage. Shift-class decomposition of the two modules (Supplementary Figure 3A, C) assigned the metabolic terms overwhelmingly to the reversed up class and the proteostasis terms to the reversed down class, with only minor progressive or stable contributions, confirming that the metabolism and proteostasis dissociation is driven by directional reversal rather than by monotonic drift. The same dissociation was reproduced at the KEGG pathway level (Supplementary Figure 3B, C), and the Functional Rewiring Index ranked biosynthesis (0.47), energy (0.36) and proteostasis (0.34) as the most extensively rewired modules (Supplementary Figure 3D). This temporal dissociation is mechanistically notable because energy supply and protein-quality control are normally coupled, proteostatic machinery is among the most ATP-demanding in the cell, and impaired bioenergetics compromises chaperone and degradative capacity (Soares et al., 2019) (Supplementary Figure 3 and Supplementary Table S3 and S4).

### 3.5 Subunit-level analysis reveals stage-dependent remodeling of respiratory-chain protein abundance

Because energy metabolism emerged as the dominant late-stage program, we dissected the mitochondrial and bioenergetic proteome in detail (Figure 4). A protein-count-weighted balance analysis across four opposition axes (Anabolic↔Catabolic, Energy↔Biosynthesis, Mitochondria↔Cytosolic, OXPHOS↔Glycolysis) showed a sign change between 72 h and 144 h, with the energy, mitochondrial and OXPHOS arms shifting most strongly toward the mutant-up direction at the later time point (Figure 4A; n = 95/76, 68/90, 58/51 and 59/16 proteins per axis pole; Supplementary Table S4). As a class, MitoCarta-3.0 proteins shifted toward the mutant-up direction at 144 h relative to non-mitochondrial proteins (Supplementary Figure 4A), and this shift was most pronounced for inner-membrane and matrix proteins (Supplementary Figure 4B). Restricting to MitoCarta-annotated DEPs, temporal classification mirrored the global pattern; 191 were late-specific, 36 early-specific and 87 shared, and among shared mitochondrial proteins reversal again predominated (n = 66) over progressive (up n = 8, down n = 6) and stable (up n = 4, down n = 3) classes, with the strongest movers including UQCR10, UQCC1, MRPL3, SLC25A12, UQCRFS1 and NDUFA3 (Figure 4B, Supplementary Table S5). Resolving the OXPHOS machinery to individual complexes showed a stage-dependent pattern (Figure 4C; Supplementary Table S5). At 72 h, all significantly changed subunits of Complex IV (cytochrome c oxidase; 6 of 6, including COX4I1 and COX6C) and Complex V (ATP synthase; 5 of 5, including ATP5F1B and ATP5F1C) were reduced, whereas Complex I and III were largely unchanged (mean log₂FC −0.01 and 0.09). At 144 h, subunits of all five complexes were broadly increased, with the largest mean increases in Complex III (mean log₂FC 0.51) and Complex I (0.36; e.g. NDUFA10, NDUFB4, NDUFB5) and smaller increases in Complex IV (0.31) and Complex V (0.24). A subset of distal subunits remained reduced at 144 h (COX5B, ATP5F1A, ATP5F1B). Reduced cytochrome c oxidase activity has been reported in HD striatum and models (Kim et al., 2010; Pickrell et al., 2011); however, whether the abundance pattern observed here alters complex assembly or respiratory activity was not tested. Glycolytic enzymes were largely unchanged or modestly reduced (Figure 4D); because protein abundance does not report flux, this does not exclude compensatory changes in glycolytic activity. Mitochondrial redox and ROS-handling proteins were dominated by late-specific changes with a smaller reversed component (Supplementary Figure 4D). Summarizing member-protein abundance into pathway scores (mean log₂FC of member proteins) showed increases for ATP synthesis, OXPHOS, fatty acid oxidation and the TCA cycle from 72 h to 144 h, and decreases for glycolysis, autophagy and the proteasome (Figure 4E), placing the 144 h state in the OXPHOS-high/glycolysis-low quadrant of the temporal energy map (Supplementary Figure 4E). These scores summarize protein abundance and do not measure pathway activity. Because the complexes differed in the size and timing of their changes, the aggregate OXPHOS score alone does not capture the subunit-level pattern (Supplementary Figure 4C; Supplementary Table S5).

**Figure 4.**
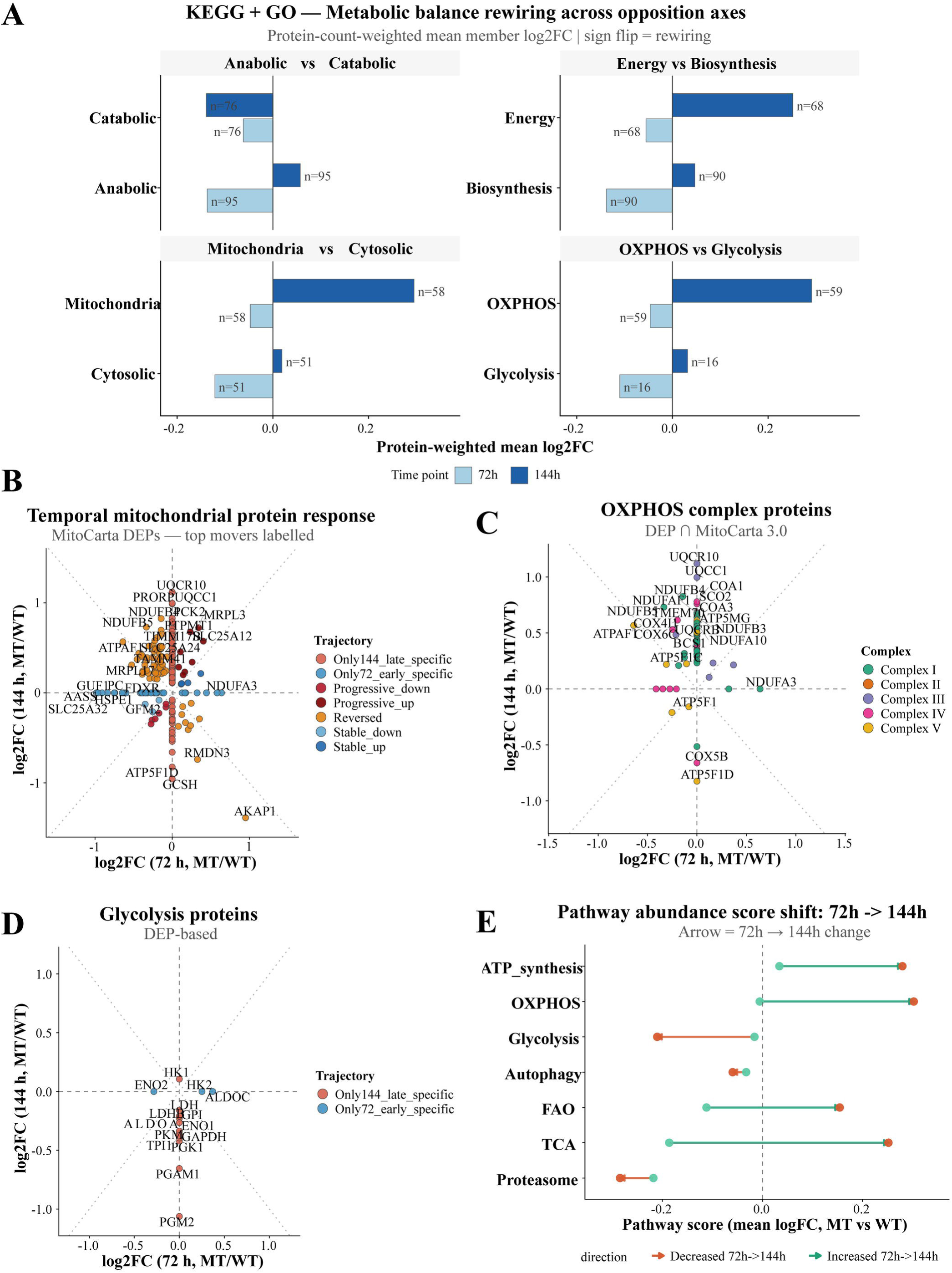
Stage-dependent remodeling of mitochondrial and respiratory-chain protein abundance. (**A**) Protein-count-weighted metabolic-balance rewiring (KEGG + GO) across four opposition axes (Anabolic↔Catabolic, Energy↔Biosynthesis, Mitochondria↔Cytosolic, OXPHOS↔Glycolysis); bars show the protein-count-weighted mean member log₂FC (MT/WT) at 72 h (light blue) and 144 h (dark blue), with per-pole protein counts annotated (n = 95/76, 68/90, 58/51 and 59/16, respectively). A change in bar sign between time points indicates rewiring of that axis. (**B**) Temporal mitochondrial-protein response: log₂FC (MT/WT) at 72 h versus 144 h for MitoCarta-3.0-annotated DEPs, coloured by trajectory class (Reversed n = 66, Progressive_up n = 8, Progressive_down n = 6, Stable_up n = 4, Stable_down n = 3, Only144/late-specific n = 191, Only72/early-specific n = 36); the top movers per trajectory are labelled. (**C**) OXPHOS complex subunits (DEP ∩ MitoCarta 3.0) plotted as log₂FC at 72 h versus 144 h and coloured by respiratory complex (I-V). Complex IV and ATP-synthase (Complex V) subunits are reduced at 72 h; at 144 h subunits of all complexes are predominantly increased, most strongly for Complex I and III. (**D**) Glycolytic enzymes (DEP-based), coloured by trajectory class; glycolytic enzymes are largely unchanged or modestly reduced (protein abundance does not report glycolytic flux). (E) Pathway-abundance score shift from 72 h (green) to 144 h (red) for ATP synthesis, OXPHOS, glycolysis, autophagy, fatty-acid oxidation (FAO), the TCA cycle and the proteasome; each pathway score is the mean member log₂FC (MT vs WT) and the arrow direction indicates whether the pathway increases (green) or decreases (red) between time points.

Because changes in respiratory-chain proteins can accompany altered mitochondrial dynamics, we examined fusion and fission proteins (Supplementary Table S2). MFN2 was quantified only at 72 h and was unchanged (log₂FC 0.03, p = 0.84); MFN1 was not quantified. The inner-membrane fusion protein OPA1 was unchanged at 72 h and increased at 144 h (log₂FC 0.30, p < 0.001), whereas the fission GTPase DNM1L/DRP1 was increased at 72 h (0.24, p = 0.015) and decreased at 144 h (−0.36, p = 0.002); FIS1 was unchanged at both time points, and the OPA1-processing protease YME1L1 followed the same pattern as OPA1 (72 h: −0.24, p = 0.005; 144 h: 0.46, p = 0.003). This pattern is compatible with a shift toward fusion at the late stage, but mitochondrial morphology and fusion were not measured; these abundance changes alone do not establish altered mitochondrial dynamics and will need to be tested in future studies.

### 3.6 Direction-aware target nomination identifies stage-matched repurposing candidates

By employing our AI-assisted direction-aware pipeline, we screened 1,711 of the 1,712 DEPs (one late-specific protein lacked a mappable target identifier and was excluded) against established resources. This approach retained 498 (29%) proteins druggable by at least one resource, of which 109 had at least one agent whose mechanism was concordant with the required correction; for 89 of these 109 proteins, at least one concordant agent was listed as approved in the source databases (Figure 5A). These counts refer to target proteins, not to individual drugs. Because approval was recorded at the target level, we also checked the current regulatory status of each candidate agent: 56 of the 89 targets had at least one currently approved concordant agent (55 with full approval and one with regional approval only), whereas the remaining 33, mostly ribosomal proteins, mapped only to agents that are not currently approved, including ataluren, whose conditional EU authorisation expired in March 2025 (Supplementary Table S6). To prioritize these candidates by interpretive tractability, we stratified all druggable DEPs into three tiers as druggable targets on a monotonic progressive trajectory (Tier 1, n = 17), druggable targets on a stable trajectory (Tier 2, n = 10), and druggable targets on a biphasic or single-time-point trajectory collapse, rebound or time-point-specific (Tier 3, n = 471) (Supplementary Figure 5A). Although Tier 3 contained the majority of druggable proteins and the largest absolute number of approved-drug targets (361 of 471 target proteins with at least one approved agent; Supplementary Table S6), its members follow transient or late-emerging trajectories whose direction of correction is intrinsically harder to interpret. The combined list of targets for Tier 1 and Tier 2 (27), by contrast, form a compact shortlist in which each candidate’s required correction (RESTORE for depleted targets, SUPPRESS for elevated targets) and mechanism concordance can be read more directly (Supplementary Figure 5C). Candidates were assigned to functional classes and organized into two layers (Figure 5B). Layer 1 comprised target-anchored agents acting on a specific druggable DEP where the mitochondrial arm converged strongly on Complex I. Because Complex I/III subunits were paradoxically elevated in mutant cells (Figure 4C), the direction-aware approach nominated agents that attenuate Complex I which together accounted for the great majority of concordant mitochondrial-target links (the approved drug metformin/metformin hydrochloride, the investigational compound ME-344 and the preclinical compound NV-128), each mapping to NDUF family and associated Complex I subunits (Supplementary Figure 5B). Layer 2 comprises pathway-level mitochondrial agents that act on bioenergetic or redox processes rather than on a single protein target. SS-31 (elamipretide; granted FDA accelerated approval for Barth syndrome in 2025, not approved for HD), MitoQ (a dietary supplement evaluated in clinical trials, not an approved drug) and idebenone (approved in the EU for Leber hereditary optic neuropathy) were included on the basis of curated HD and energy-metabolism literature and positioned against the whole Complex I/OXPHOS node in the drug-target network (Figure 5B). The Complex I-associated targets (26, Figure 5B) that anchor this network are supported by database evidence and are direction-consistent; each is druggable by all three resources and lies on a biphasic or late trajectory (Tier 3). The SUPPRESS-concordant action resolved the two single-time-point subunits that were invisible on the biphasic scatter (72 h to 144 h temporal profiles, Supplementary Figure 5D).

**Figure 5.**
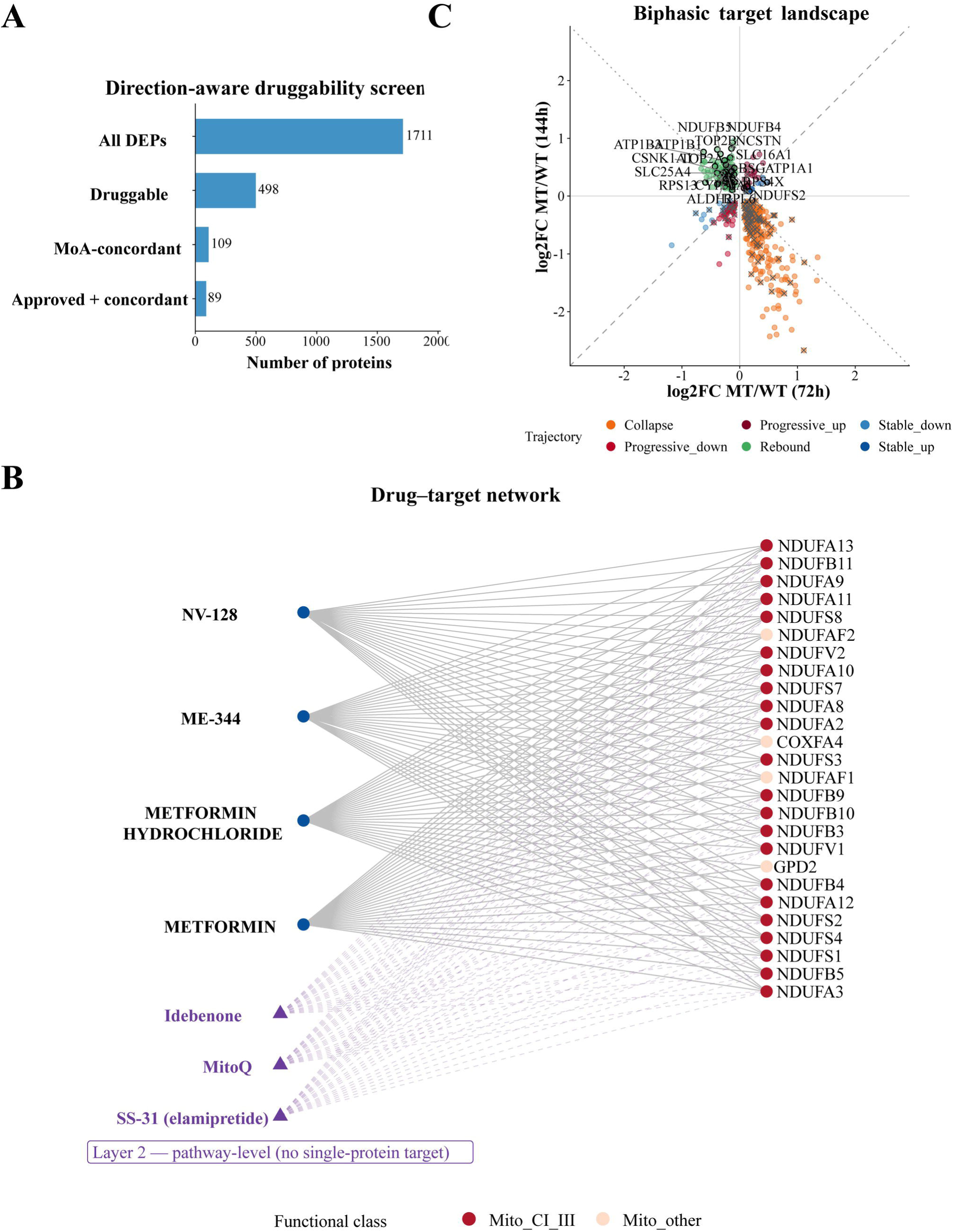
Direction-aware drug repurposing identifies stage-matched candidates and a putative early intervention window. (**A**) Direction-aware druggability screening funnel: all DEPs with a mappable target identifier (n = 1,711) → druggable by at least one resource (n = 498) → mechanism-of-action-concordant with the required correction (n = 109) → approved and concordant (n = 89; approval as recorded in the source databases, of which 56 targets have a currently approved concordant agent). (**B**) Two-layer drug-target network. Layer 1 (solid edges) ,comprises target-anchored hubs — metformin and metformin hydrochloride (approved), ME-344 (investigational) and NV-128 (preclinical), converging on Complex I (NDUF family) and associated subunits; target nodes are coloured by functional class (Mito_CI_III, red; Mito_other, grey). Layer 2 (dashed edges, triangles) comprises pathway-level mitochondrial agents — SS-31 (elamipretide; approved for Barth syndrome), MitoQ (dietary supplement) and idebenone (approved in the EU for LHON) — that act on the bioenergetic/redox node rather than on a single protein target. (C) Biphasic target landscape: DEPs shared between the two time points (n = 519) plotted by log₂FC (MT/WT) at 72 h versus 144 h and coloured by trajectory class (Collapse, Rebound, Progressive_up, Progressive_down, Stable_up, Stable_down). Black-ringed points are druggable and mechanism-concordant (actionable) and grey crosses are druggable but mechanism-discordant (wrong-direction). Collapse, increased at 72 h and decreased at 144 h; Rebound, decreased at 72 h and increased at 144 h. Among proteins shared between the two time points, concordant candidates are mostly in the rebound class (29 of 33), and no collapse protein has a concordant agent.

Beyond the mitochondrial node, the direction-aware screen recovered actionable candidates for the proteostasis and RNA-regulatory arms (Supplementary Figure 5E). Ribosomal and proteasomal proteins that were progressively depleted or reversed nominated translation-modulating agents (none of which is currently approved; Supplementary Table S6) and proteasome-directed agents, while individual high-confidence targets included IMPDH2 (nominating the approved immunomodulators mycophenolate and mizoribine), the RNA-binding protein FUS (tretinoin), and the DNA mismatch-repair factor MSH3 an established genetic modifier of HD somatic instability nominating antimetabolite chemistry. Two agents, olesoxime and triheptanoin, are of particular interest because both have prior neurodegeneration/energy-metabolism rationale (Mochel, 2017; Weber et al., 2019). Finally, plotting the DEPs shared between the two time points on the 72 h vs 144 h landscape (Figure 5C) and the temporal profile of the mitochondrial targets (Supplementary Figure 5D) showed that almost all concordant targets were proteins elevated in mutant cells for which an inhibitor was available (108 of 109; Supplementary Table S6): late-specific proteins increased at 144 h (n = 57), rebound proteins (decreased at 72 h and increased at 144 h; n = 29), early-specific proteins increased at 72 h (n = 18) and a few progressive or stable increased proteins (n = 4), with a single concordant activator for a depleted target. None of the 270 collapse proteins (increased at 72 h and decreased at 144 h) had a concordant agent, because their correction would require restoring a depleted protein, whereas the available agents were inhibitors. Relative to their share of all DEPs, rebound proteins were over-represented among concordant targets (27% vs 10%). This distribution therefore reflects the direction of change and the mechanisms of the available agents, and does not by itself indicate at which stage intervention would be most effective, which would require intervention experiments.

## 4 Discussion

Our results provided a temporally resolved map of proteome remodeling driven by polyglutamine-expanded huntingtin, and its central finding was that mutant huntingtin did not simply amplify a fixed set of perturbations over time but was instead associated with a two-stage trajectory in which the majority of shared proteins reversed the direction of their abundance change between an early and a late time point (Figure 3B, C). The early proteome (72 h) was characterized by increased abundance of RNA-splicing, ribonucleoprotein-biogenesis and translational proteins, a pattern that may reflect an adaptive attempt to preserve gene-expression fidelity under accumulating mutant-protein stress. The disrupted RNA metabolism and RNA-binding-protein homeostasis are increasingly recognized pathogenic axes in HD and related neurodegeneration (Picchiarelli and Dupuis, 2020; Matera and Wang, 2014; Aviner et al., 2024). By the late time point (144 h), the proteome changes indicated broad protein depletion and to a dominant energy-metabolism signature, consistent with the mitochondrial dysfunction and translational failure repeatedly documented in HD tissue and models (Mochel and Haller, 2011; Eshraghi et al., 2021; Kim et al., 2010; Lisowski et al., 2024).

The scale of directional reversal of shared DEPs (85%; 95% at FDR < 0.05) suggests that, in this model, the mutant proteome is reorganized between the two time points rather than progressively intensified (Figure 3B). We emphasize that this transition is restricted to proteins that remain significant at both time points as proteins that were perturbed early and then normalized are captured in the early-specific class. Both the transition and the early-specific populations describe transient early responses that are not sustained, and the two together constitute the majority of early changes. The convergence of enrichment analyses early spliceosome and nucleocytoplasmic-transport signatures giving way to late oxidative phosphorylation and respiratory chain signatures are consistent with a sequence in which an early RNA-regulatory response precedes late remodeling of energy-metabolism proteins (Figure 3D); with two time points, the intermediate steps of this sequence cannot be resolved.

A further observation is that respiratory-chain proteins changed differently between complexes and between time points. Complex IV and ATP synthase subunits were reduced early, whereas at 144 h subunits of all complexes increased, with larger increases in Complex I and III than in Complex IV and V (Figure 4C). An aggregate OXPHOS score therefore conceals this subunit-level heterogeneity. An uneven change in the relative abundance of respiratory-chain components could affect complex assembly or electron flow, and reduced cytochrome c oxidase activity has been reported in HD tissue and models (Mochel and Haller, 2011; Pickrell et al., 2011). Although protein abundance does not report complex assembly or respiratory activity, we think that this pattern may contribute to a developing energy failure in mutant cells; functional measurements, such as oxygen consumption, Complex IV activity and cellular ATP, are needed to test this possibility. The dissociation we observe between the metabolic and proteostatic programs further indicates that these two systems change on different schedules an important point given that the ubiquitin-proteasome system, autophagy, and chaperone networks are themselves progressively compromised in HD and depend on an intact energy supply (Soares et al., 2019; Ortega et al., 2007; Sarkar and Rubinsztein, 2008; Ortega and Lucas, 2014; Harding and Tong, 2018).

Taken together, the data supports a model in which early disruption of RNA regulatory networks and compensatory activation of translational machinery preceded late remodeling of mitochondrial energy-metabolism and proteostasis proteins. This two-stage pattern suggests a putative early window, to be tested in preclinical intervention studies, in which interventions aimed at RNA quality control, nucleocytoplasmic transport or proteostatic support might delay the late mitochondrial changes and a possible progression towards energy failure (Ross and Tabrizi, 2011; Tabrizi et al., 2020). Whether the late changes can still be reversed was not tested here. The direction-aware repurposing analysis reported here provides putative candidates for testing this idea: agents that attenuate Complex I for the respiratory-chain imbalance and translation or proteostasis modulators for the early RNA-regulatory changes. These are computational nominations that will need experimental validation.

Bioenergetic trials have repeatedly targeted the respiratory chain without regard to stage or direction. High-dose coenzyme Q10 (with remacemide) in the CARE-HD trial and creatine in CARE-HD-era studies and the large CREST-E trial showed no slowing of clinical decline (Huntington Study Group, 2001; Hersch et al., 2017; Verbessem et al., 2003). Antioxidants targeting mitochondria such as MitoQ and the cardiolipin-stabilizing peptide SS-31/elamipretide have not translated into approved HD therapies despite a strong mechanistic rationale and evidence that they can delay disease in neurodegeneration models (Yin et al., 2016; Faô and Rego, 2021; Tung et al., 2025). Metformin, the Complex I targeting agent, has independent epidemiological and preclinical support in HD and is under active consideration as a repurposing candidate as in our analyses (Trujillo-Del Río et al., 2022). Approaches that raise rather than blunt mitochondrial capacity PGC-1α/biogenesis activators such as bezafibrate (Johri et al., 2012; Chandra et al., 2016), the anaplerotic substrate triheptanoin (Weber et al., 2019), the mitochondrial-permeability-transition modulator olesoxime (Mochel, 2017), and the transglutaminase-linked agent cystamine (Borrell-Pagès et al., 2006) have likewise been explored with limited clinical success. Our data may offer one possible explanation for why these energy-focused efforts underperformed (Figure 4C, E). In our model, the late-stage OXPHOS signature was not a uniform loss but an uneven increase, with larger rises in proximal Complex I/III subunits than in distal Complex IV/V subunits, following an early reduction of distal subunits. An intervention that indiscriminately stimulates the respiratory chain could therefore reinforce the proximal imbalance we observe, whereas the direction-aware approach applied here nominates Complex I-attenuating agents. Because the Complex I subunits targeted by these agents were increased at the late time point, the stage at which such treatment would be most appropriate is not defined by the present data and needs to be tested experimentally. This suggests that mitochondrial repurposing in HD might be considered not only as ‘boosting energy metabolism’ but also as restoring a stage-appropriate direction of change, and it provides testable drug–target pairings (Figure 5B). Complex I attenuation and proteostasis support would represent a different strategy from the broad antioxidant and bioenergetic approaches of prior trials; whether and when such treatment is beneficial remains to be tested experimentally. Beyond the mitochondrial axis, the recovery of MSH3 whose suppression lowers somatic CAG expansion is an emerging disease-modifying strategy (Bunting et al., 2025; GeM-HD Consortium, 2025). It offers a druggable, direction-concordant target that illustrates that the same unbiased map mechanistically orthogonal opportunities alongside the energy-metabolism core. Framing these nominations by tier helps to prioritize them. The largest block of druggable, high-confidence targets including the entire Complex I network that dominates the drug-target map falls into Tier 3, because these subunits change only at the late time point or follow rebound trajectories so that even well-supported agents such as metformin act on a stage at which the required direction of correction is least clear (Supplementary Figure 5A). The direction-resolved Tier 1/Tier 2 shortlist instead isolates the 27 targets whose progressive or stable behavior makes their required correction unambiguous, and it is here rather than in the numerically dominant but temporally transient Tier 3 that early-window intervention is most defensible for future testing (Supplementary Figure 5C and Supplementary Table S6). Several depleted proteostasis proteins would, in principle, require restoration. For instance, the proteasome subunits (PSMC2 and PSMD4) and the purine-biosynthesis enzyme (IMPDH2) matched only to inhibitory chemistry and were therefore flagged mechanism-discordant, so their inclusion marked a target worth pursuing rather than a ready-to-use agent (Supplementary Figure 5C).

The design of this study was chosen to answer a specific question: how the proteome, and in particular energy metabolism, changes as polyQ aggregates form. HEK293T cells were used because they allow efficient, synchronized expression and reproducible aggregation of polyQ-expanded huntingtin within a defined time frame, which time-resolved proteomics requires, and because they express several neuronal-lineage proteins (Shaw et al., 2002). The findings therefore describe aggregation-associated, cell-autonomous changes rather than neuron-specific metabolism. The two time points were selected to capture an early stage at which aggregates have formed and a later stage with a lower per-cell aggregate load (Supplementary Figure 1), and the proteomic data report protein abundance. The respiratory-chain and drug-repurposing results should therefore be regarded as hypotheses generated from protein abundance data. Building on this map, future studies will need to measure bioenergetic function directly (oxygen consumption, Complex IV activity and ATP levels), validate selected changes in endogenous proteins, examine mitochondrial morphology and dynamics, test the nominated candidates with treatments applied at early and late stages, and examine whether the same temporal pattern occurs in HD neuronal models.

## 5 Conclusion

Using quantitative global proteomics in a HEK293T model, we describe a two-stage proteomic response to polyglutamine-expanded huntingtin: an early increase in RNA-regulatory and translational proteins at 72 h, followed by depletion of many of these proteins and remodeling of mitochondrial energy-metabolism proteins at 144 h, with 85% of shared differentially abundant proteins changing direction between time points. Subunit-level analysis showed that respiratory-chain complexes changed unevenly, with an early reduction of Complex IV and ATP synthase subunits and a later increase that was largest for Complex I and III. These abundance patterns suggest that mutant cells may be progressing towards energy failure, a possibility that remains to be confirmed with functional measurements. The time-resolved map supports a stage-matched, direction-aware approach to drug repurposing and nominates candidates such as the Complex I inhibitor metformin, whose neuroprotective rationale has been reviewed previously (Rotermund et al., 2018; Du et al., 2022). Future work should test these candidates in neuronal and in vivo HD models, with functional readouts such as oxygen consumption and ATP levels and with treatments applied at different stages, to determine whether a putative early window for intervention exists.

## Declarations

## Acknowledgements

The authors thank the Cord Omics team for their support, especially Simay Akpinar and İbrahim Yazgan for technical assistance, and Prof. Dr. Işıl Aksan Kurnaz for her valuable comments and support. Computational analyses were performed with the assistance of an autonomous large-language-model research agent (Claude Code and Claude Science, Anthropic) (Anthropic, 2026). All drug–target associations were drawn from established public resources under an explicit constraint, and all AI-assisted outputs were reviewed and verified by the authors. The complete prompts used for the AI-assisted target-nomination step are embedded in the analysis scripts deposited in the code repository (see Data Availability Statement). A preprint version of this manuscript has been submitted to bioRxiv (doi: 10.64898/2026.08.21.746221; Sonmez et al., 2026).

## Data availability

The mass-spectrometry proteomics data have been deposited to the ProteomeXchange Consortium via the PRIDE partner repository under accession PXD082949. The processed differential-expression tables, temporal-classification results and the annotated drug-target table are provided in the Supplementary Material. All analysis scripts including FragPipe/IonQuant search and quantification parameters, the R workflow for NormalyzerDE normalization, missing-value imputation, and DEqMS differential expression analysis, clusterProfiler functional enrichment, and the drug-repurposing target-nomination pipeline are publicly available at https://github.com/cimenlab/Temporal-Proteomics-of-mutant-HTT-Biphasic-Molecular-Cascade-KAP-.

## Author contributions

HC and GA conceptualized the study. ES, MS, GA, MK, and HC designed the methodology. ES, PM, CO, MS, GA, MK, and HC performed the investigation. ES, CO, and HC carried out formal analysis. GA, MK, and HC contributed resources. ES, PM, and CO prepared visualization. ES and HC wrote the original draft. ES, PM, MS, GA, MK, and HC reviewed and edited the manuscript. GA and HC supervised the project and acquired funding. All authors read and approved the final manuscript.

## Supplementary Material

### Supplementary Tables

**Table S1. Quantified proteins and quality-control summary.** MaxLFQ protein-level intensities (FragPipe/IonQuant) at 72 h and 144 h, proteotypic-peptide counts and removed contaminant entries. Sheets: Quantified_72h_sheet1, Quantified_144h_sheet1, Proteotypic_Sheet_1, Contaminants_removed_Sheet_1 (4 data sheets). All values were generated with FragPipe/IonQuant (MaxLFQ) and DEqMS; column headers follow the Proteome Discoverer naming convention used by the analysis scripts, and all ratios are expressed as MT/WT.

**Table S2. Master differentially abundant protein table (72 h and 144 h).** One row per protein with log₂ fold-change (MT/WT) and DEqMS p-value at both time points, temporal class (early-specific / late-specific / persistent) and progression class (progressive / stable / reversed), together with the per-time-point DEP lists, the cross-time-point overlap and group sizes. DEqMS p < 0.05. Sheets: MASTER_DEP_table, DEP_72h_DEPs, DEP_144h_DEPs, Overlap_Counts, Overlap_Shared_DEPs, Overlap_Only_72h, Overlap_Only_144h, TimeProg_Summary, Group_sizes_Sheet_1, Top_labeled_genes_Sheet_1, Class_counts. Revision sheets (prefix REV_): FDR_overlap_reversal, FDR_classes_per_protein, Interaction_summary, Interaction_model, Sensitivity_no_imputation, Sensitivity_count_var, Imputation_in_DEPs, Imputation_flags_72h, Imputation_flags_144h, Mito_dynamics (21 data sheets). Benjamini–Hochberg-adjusted p-values (padj) are given in DEP_72h_DEPs and DEP_144h_DEPs.

**Table S3. Functional over-representation (GO:BP and KEGG).** clusterProfiler over-representation results for the up- and down-regulated sets at each time point, for the temporal classes (Only72 / Shared / Only144) and for the progression classes, together with the metabolism-versus-proteostasis module split (genome-wide annotation background). Revision sheets (prefix REV_) contain the sensitivity analysis with FDR-filtered DEP lists and all quantified proteins as background (Supplementary Figure 6). Term membership was drawn from database annotation rather than from the enrichment output. 18 data sheets.

**Table S4. Metabolic rewiring: opposition-axis membership, balance scores and rewiring index.** Four opposition axes (Anabolic↔Catabolic, Energy↔Biosynthesis, Mitochondria↔Cytosolic, OXPHOS↔Glycolysis) with GO:BP and KEGG term and protein membership and protein-count-weighted mean member log₂FC (MT/WT) per pole at 72 h and 144 h; the metabolism-versus-proteostasis dissociation tables and shift-class assignments; the Metabolic Rewiring Index (MRI) and the Functional Rewiring Index (FRI, KEGG + GO, overall and by progression class); and the protein-level rewiring and reversal sets. 19 data sheets.

**Table S5. Mitochondrial and bioenergetic compendium (MitoCarta 3.0).** MitoCarta-annotated DEPs with trajectory class; sub-organellar localization (inner membrane, matrix, intermembrane space, outer membrane); OXPHOS complex membership (I–V) with per-complex summaries based on DEqMS statistics (log2FC, p and BH-adjusted p); ROS/redox markers; glycolytic enzymes; pathway abundance scores for the 72 h → 144 h transition; and complex and energy-balance scores. 15 data sheets.

**Table S6. Direction-aware drug-repurposing target table.** Per-DEP druggability annotation (DGIdb, Open Targets, ChEMBL, DrugBank) with evidence count, candidate agents, mechanism of action and highest clinical phase, the required correction (RESTORE for depleted targets, SUPPRESS for elevated targets), mechanism concordance, Layer 1 versus Layer 2 assignment, priority tier (Tier 1–3) and the regulatory status of individual named agents. For mechanism-concordant targets with an approved agent in the source databases, the candidate agents that are currently approved are listed, and the current status of each agent is given in a separate sheet (REV_Agent_status_lookup). Drug–target associations were taken only from verified database records under a no-invented-links constraint. 7 data sheets.

*Each supplementary table is provided as a single multi-sheet Excel workbook containing a README sheet and a sheet index*.

**Supplementary Figure 1. Aggregation and construct expression in the huntingtin cell model at the proteomic time points.** (A) Representative 10× fluorescence images of Htt-Q23 and Htt-Q74 cells at 72 h and 144 h post-transfection (GFP-Htt, green; nuclei, blue). (B) Representative 40× images showing a diffuse cytoplasmic Htt-Q23 signal and discrete puncta in Htt-Q74 cells. (C) Aggregation kinetics of Htt-Q74 cells across 10× fields, normalized to 72 h: integrated GFP density and mean cell count (bars, left axis) and aggregate load per cell (line, right axis). (D) Morphology of the segmented GFP signal in Htt-Q23 and Htt-Q74 cells at 72 h and 144 h: mean signal area (px², bars) and circularity index (0–1, diamonds); the Q23 signal corresponds to whole cell bodies and the Q74 signal to individual puncta. Mean ± SD of n ≥ 18 fields per group from 3 independent transfections; two-tailed unpaired Student’s t-test (C, 144 h vs 72 h; D, Q23 vs Q74 at each time point); ** p < 0.01, *** p < 0.001. Scale bars: 138.8 µm (A), 34.7 µm (B). (E) Anti-GFP immunoblot of whole-cell lysates from GFP, Q23 and Q74 cells at 72 h and 144 h, with β-actin as loading control; the GFP-Htt fusions (Q23, Q74) migrate above free GFP. (F) Total-protein reference stain of the same samples with molecular-weight markers (15-170 kDa), confirming comparable loading across conditions and time points.

**Supplementary Figure 2. Extended functional over-representation atlas (GO:BP and KEGG).** (**A**) GO:BP over-representation across the up- and down-regulated DEP sets at each time point (72h_UP, 72h_DOWN, 144h_UP, 144h_DOWN). (**B**) KEGG over-representation across the same four sets. (**C**) GO:BP over-representation across the temporal classes (Only72, Shared, Only144). (**D**) KEGG over-representation across the temporal classes. Throughout, dot size = gene ratio, colour = adjusted p-value and the number of mapped genes per group is annotated on the x-axis.

**Supplementary Figure 3. Extended functional rewiring: shift-class decomposition, KEGG trajectory and the functional rewiring index.** (**A**) GO:BP shift-class decomposition for the Metabolism (upper) and Proteostasis (lower) modules; each term is positioned by its score at 72 h versus 144 h, coloured by shift class (Progressive_down, Reversed_down, Reversed_up, Stable_down) and sized by |Δ score|. Metabolic terms fall overwhelmingly into the reversed-up class and proteostasis terms into the reversed-down class. (B) KEGG pathway-trajectory plot (score at 72 h versus 144 h) coloured by module (Metabolism, red; Proteostasis, blue); point size = |Δ score|. (**C**) KEGG shift-class decomposition for the Metabolism (upper) and Proteostasis (lower) modules, plotted as in (A). (**D**) Functional Rewiring Index (FRI; mean |Δ score| per category, KEGG + GO) for the Biosynthesis (0.47), Energy (0.36), Proteostasis (0.34), Lipid (0.22) and Amino-acid (0.16) modules.

**Supplementary Figure 4. Extended mitochondrial analysis: sub-organellar localization, OXPHOS complex scores, redox markers and the temporal energy map.** (A) Density distribution of log₂FC (MT/WT) for MitoCarta-3.0 proteins versus non-mitochondrial proteins at 72 h and 144 h. (B) Sub-mitochondrial localization scores (MIM = inner membrane; matrix; IMS = intermembrane space; MOM = outer membrane; membrane; matrix|IMS; unknown) at 72 h versus 144 h; dot size = number of proteins, colour = mean log₂FC. (C) OXPHOS complex scores (Complexes I-V) at 72 h versus 144 h; dot size = number of proteins, colour = mean log₂FC. Complex I carries the largest protein count; at 144 h all complexes show a positive mean log₂FC, highest for Complex III. (D) ROS/redox markers (DEPs intersecting MitoCarta redox pathways): log₂FC at 72 h versus 144 h, coloured by trajectory (Only144/late-specific, Only72/early-specific, Reversed). (E) Temporal energy map: OXPHOS score versus glycolysis score (mean member log₂FC, MT vs WT) at 72 h (green) and 144 h (orange), with the arrow indicating the direction of the transition. The 144 h state moves into the OXPHOS-high/glycolysis-low quadrant.

**Supplementary Figure 5. Extended direction-aware druggability screen: priority tiers, drug convergence and actionable candidates.** (**A**) Druggable targets across the biphasic landscape: every DEP plotted by log₂FC (MT/WT) at 72 h versus 144 h and coloured by priority tier over the non-druggable background (grey, n = 1,213); Tier 1 (monotonic progressive trajectory, n = 17), Tier 2 (stable trajectory, n = 10) and Tier 3 (biphasic or single-time-point trajectory, n = 471). The number of Tier-3 targets that map to agents listed as approved in the source databases (361 of 471) is reported in Supplementary Table S6. (**B**) Drug convergence on mitochondrial targets (top 15 agents by number of concordant mitochondrial targets): metformin and metformin hydrochloride (26 targets each), ME-344 and NV-128 (25 each) far exceed all other agents, which map to a single target. (**C**) Tier 1 and Tier 2 actionable shortlist (27 targets) ordered by log₂FC (MT/WT) at 144 h and coloured by the required correction (RESTORE, target depleted in mutant, blue; SUPPRESS, target elevated in mutant, red); filled points are mechanism-concordant and open points mechanism-discordant, with the Tier 1/Tier 2 boundary marked. (**D**) Mitochondrial druggable targets — temporal profile: log₂FC (MT/WT) at 72 h (light) and 144 h (dark) for Mito_CI_III and Mito_other targets, including single-time-point proteins (Only72/Only144) that are not visible on the biphasic scatter. (**E**) Actionable non-mitochondrial candidates (top 30): concordant, approved agents complementing the mitochondrial arm, coloured by functional class (Lipid, Other, Proteostasis, RNA_processing) and labelled with the highest-phase drug (log₂FC MT/WT, persistent).

**Supplementary Figure 6. Sensitivity analysis of functional over-representation with FDR-filtered DEP lists and a quantified-proteome background.** (A) GO:BP and (B) KEGG over-representation (clusterProfiler compareCluster) for DEP sets defined at Benjamini–Hochberg-adjusted p < 0.05 (72h_UP, n = 209; 72h_DOWN, n = 161; 144h_UP, n = 629; 144h_DOWN, n = 655) and for the temporal classes (Only72, n = 123; Shared, n = 247; Only144, n = 1,037), using all proteins quantified at either time point as background. Only sets with at least one significant term are shown: Only72 had no significant GO:BP or KEGG term, and 144h_DOWN and Only144 had no significant KEGG term. Numbers under the x-axis give the genes of each set mapped to the annotation. The top-ranked terms per set (Benjamini– Hochberg-adjusted p < 0.05) are shown; dot size = gene ratio, colour = adjusted p-value; long GO:BP term names are truncated. Complete results are given in Supplementary Table S3 (REV_ENR sheets).

