## Supplementary for "Time-Resolved Proteomics of a Polyglutamine-Expanded Huntingtin in vitro Cell Model Reveals Early RNA-Regulatory Compensation and Late Respiratory-Chain Remodeling"

### Supplementary Material

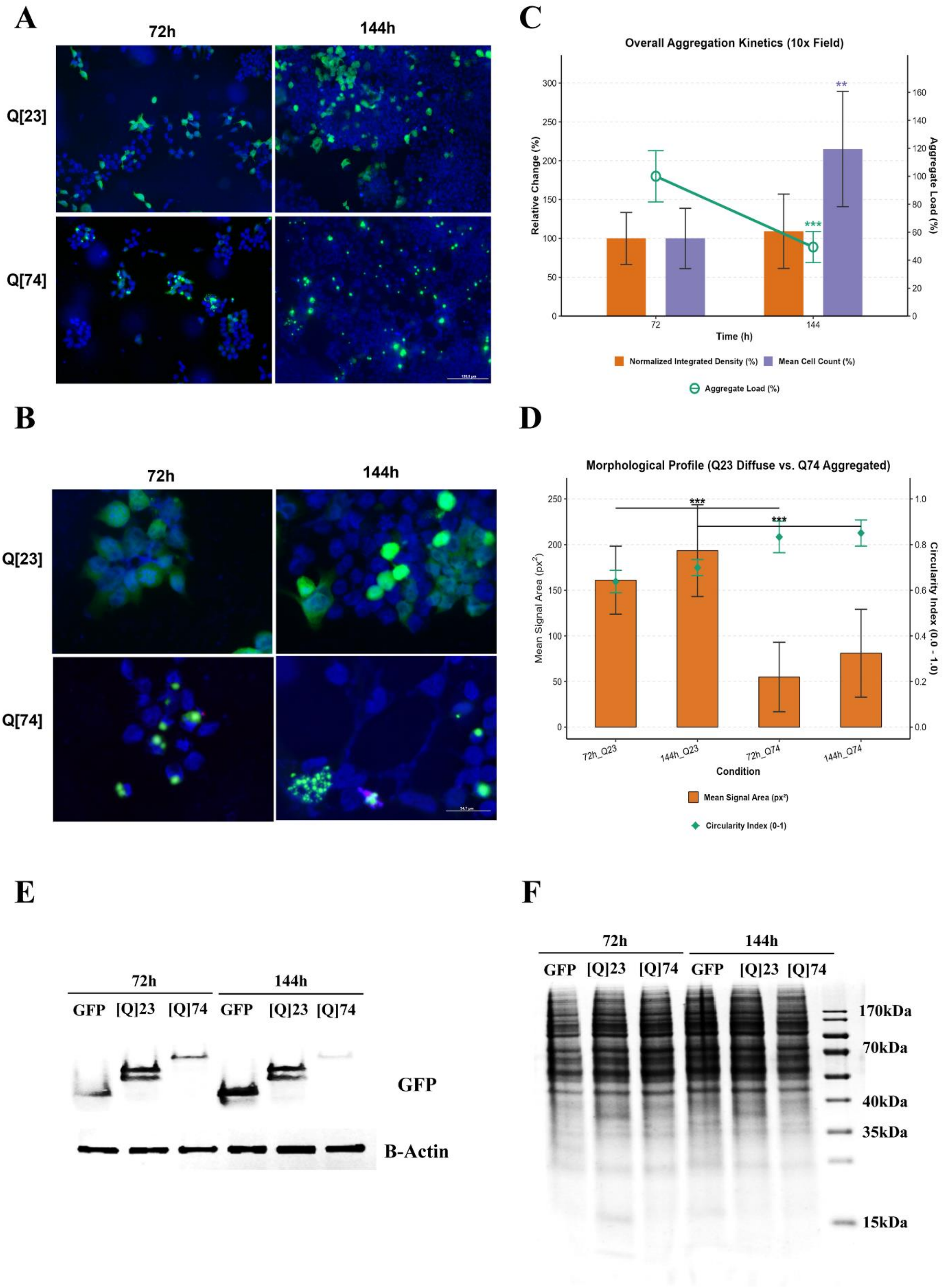


**Supplementary Figure 1.** Aggregation and construct expression in the huntingtin cell model at the proteomic time points. (A) Representative 10× fluorescence images of Htt-Q23 and Htt-

Q74 cells at 72 h and 144 h post-transfection (GFP-Htt, green; nuclei, blue). (B) Representative 40× images showing a diffuse cytoplasmic Htt-Q23 signal and discrete puncta in Htt-Q74 cells. (C) Aggregation kinetics of Htt-Q74 cells across 10× fields, normalized to 72 h: integrated GFP density and mean cell count (bars, left axis) and aggregate load per cell (line, right axis). (D) Morphology of the segmented GFP signal in Htt-Q23 and Htt-Q74 cells at 72 h and 144 h: mean signal area (px$^2$, bars) and circularity index (0–1, diamonds); the Q23 signal corresponds to whole cell bodies and the Q74 signal to individual puncta. Mean ± SD of $n \geq 18$ fields per group from 3 independent transfections; two-tailed unpaired Student's t-test (C, 144 h vs 72 h; D, Q23 vs Q74 at each time point); ** $p < 0.01$, *** $p < 0.001$. Scale bars: 138.8 µm (A), 34.7 µm (B). (E) Anti-GFP immunoblot of whole-cell lysates from GFP, Q23 and Q74 cells at 72 h and 144 h, with β-actin as loading control; the GFP-Htt fusions (Q23, Q74) migrate above free GFP. (F) Total-protein reference stain of the same samples with molecular-weight markers (15-170 kDa), confirming comparable loading across conditions and time points.

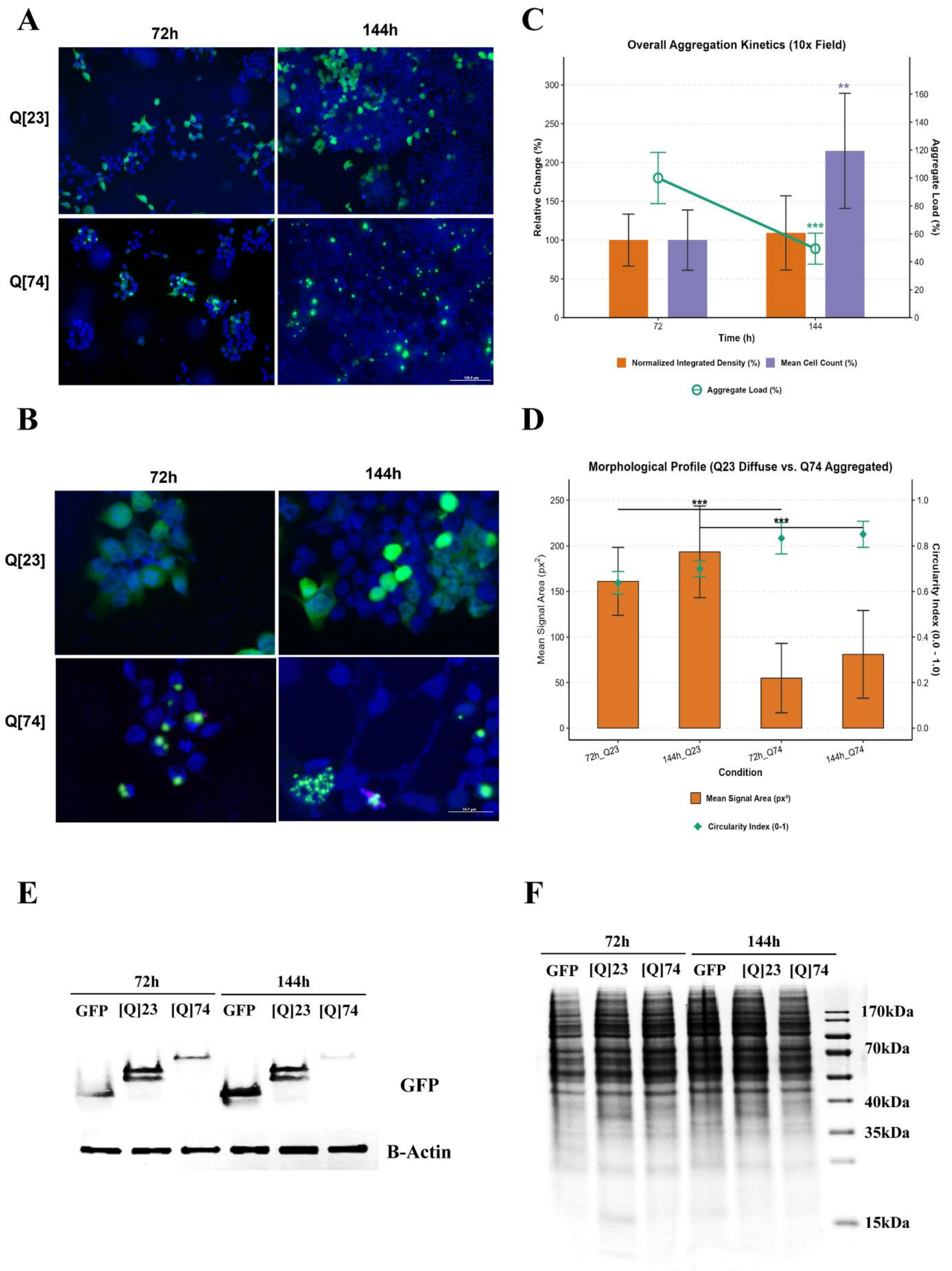


**Supplementary Figure 2. Extended functional over-representation atlas (GO:BP and KEGG).** (**A**) GO:BP over-representation across the up- and down-regulated DEP sets at each time point (72h_UP, 72h_DOWN, 144h_UP, 144h_DOWN). (**B**) KEGG over-representation across the same four sets. (**C**) GO:BP over-representation across the temporal classes (Only72, Shared, Only144). (**D**) KEGG over-representation across the temporal classes. Throughout, dot size = gene ratio, colour = adjusted p-value and the number of mapped genes per group is annotated on the x-axis.

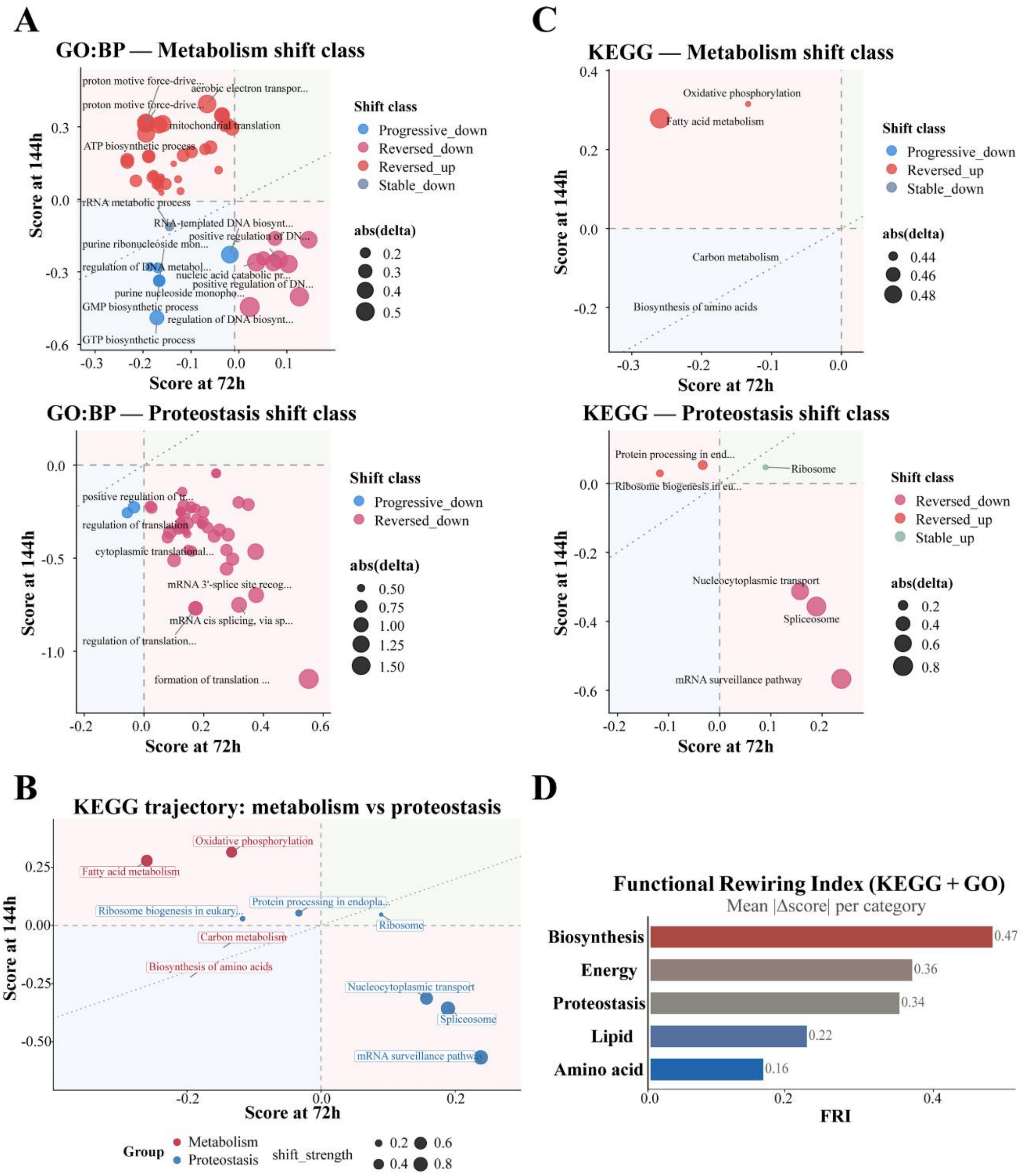


**Supplementary Figure 3. Extended functional rewiring: shift-class decomposition, KEGG trajectory and the functional rewiring index.** (**A**) GO:BP shift-class decomposition for the Metabolism (upper) and Proteostasis (lower) modules; each term is positioned by its score at 72 h versus 144 h, coloured by shift class (Progressive_down, Reversed_down, Reversed_up, Stable_down) and sized by |Δ score|. Metabolic terms fall overwhelmingly into the reversed-up class and proteostasis terms into the reversed-down class. (B) KEGG pathway-trajectory plot (score at 72 h versus 144 h) coloured by module (Metabolism, red; Proteostasis, blue); point size = |Δ score|. (**C**) KEGG shift-class decomposition for the Metabolism (upper) and Proteostasis (lower) modules, plotted as in (A). (**D**) Functional Rewiring Index (FRI; mean |Δ score| per category, KEGG + GO) for the Biosynthesis (0.47), Energy (0.36), Proteostasis (0.34), Lipid (0.22) and Amino-acid (0.16) modules.

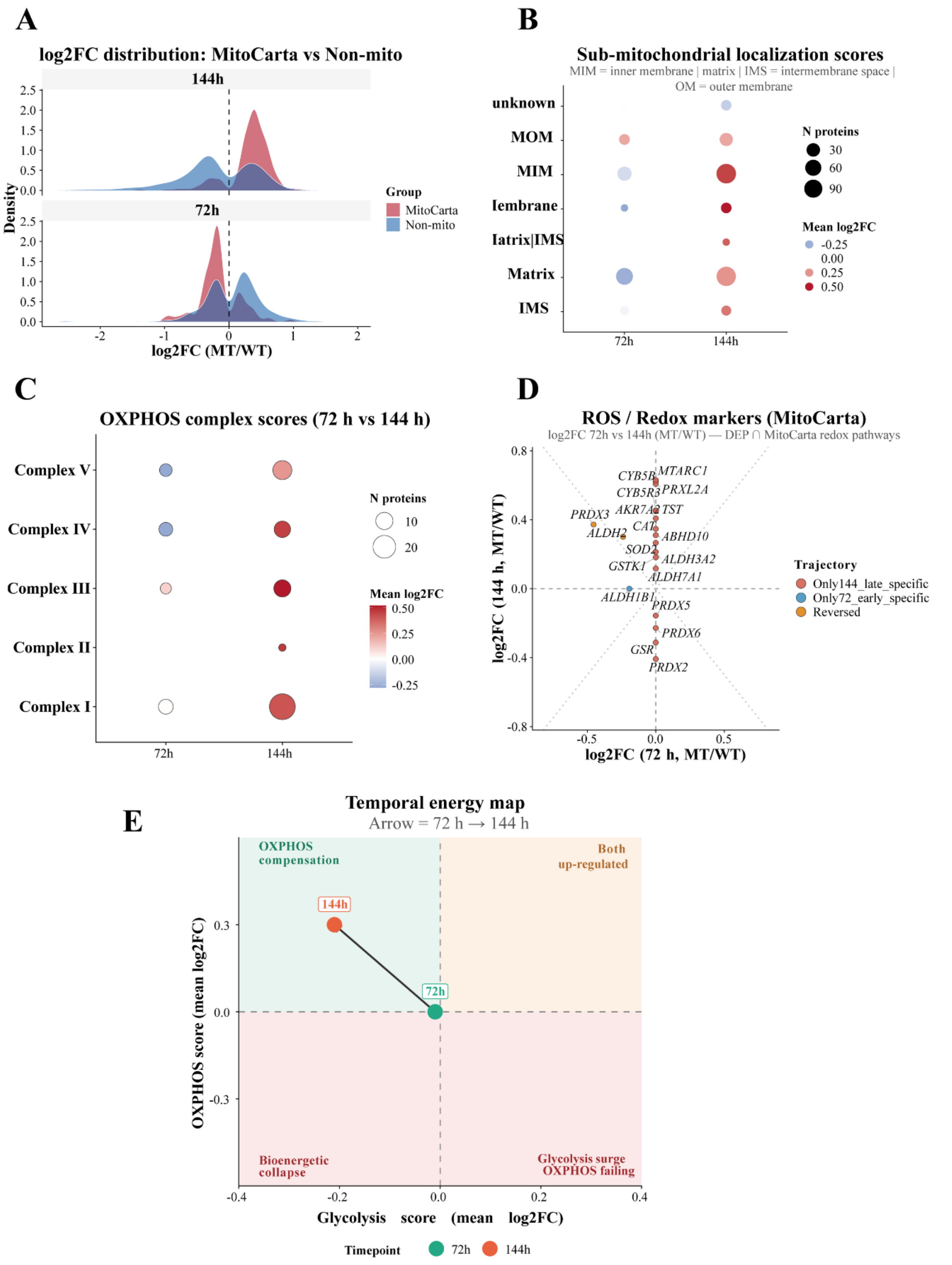


**Supplementary Figure 4. Extended mitochondrial analysis: sub-organellar localization, OXPHOS complex scores, redox markers and the temporal energy map.** (**A**) Density distribution of $log_2FC$ (MT/WT) for MitoCarta-3.0 proteins versus non-mitochondrial proteins at 72 h and 144 h. (**B**) Sub-mitochondrial localization scores (MIM = inner membrane; matrix; IMS = intermembrane space; MOM = outer membrane; membrane; matrix|IMS; unknown) at

72 h versus 144 h; dot size = number of proteins, colour = mean $log_2FC$. (**C**) OXPHOS complex scores (Complexes I-V) at 72 h versus 144 h; dot size = number of proteins, colour = mean $log_2FC$. Complex I carries the largest protein count; at 144 h all complexes show a positive mean $log_2FC$, highest for Complex III. (**D**) ROS/redox markers (DEPs intersecting MitoCarta redox pathways): $log_2FC$ at 72 h versus 144 h, coloured by trajectory (Only144/late-specific, Only72/early-specific, Reversed). (**E**) Temporal energy map: OXPHOS score versus glycolysis score (mean member $log_2FC$, MT vs WT) at 72 h (green) and 144 h (orange), with the arrow indicating the direction of the transition. The 144 h state moves into the OXPHOS-high/glycolysis-low quadrant.

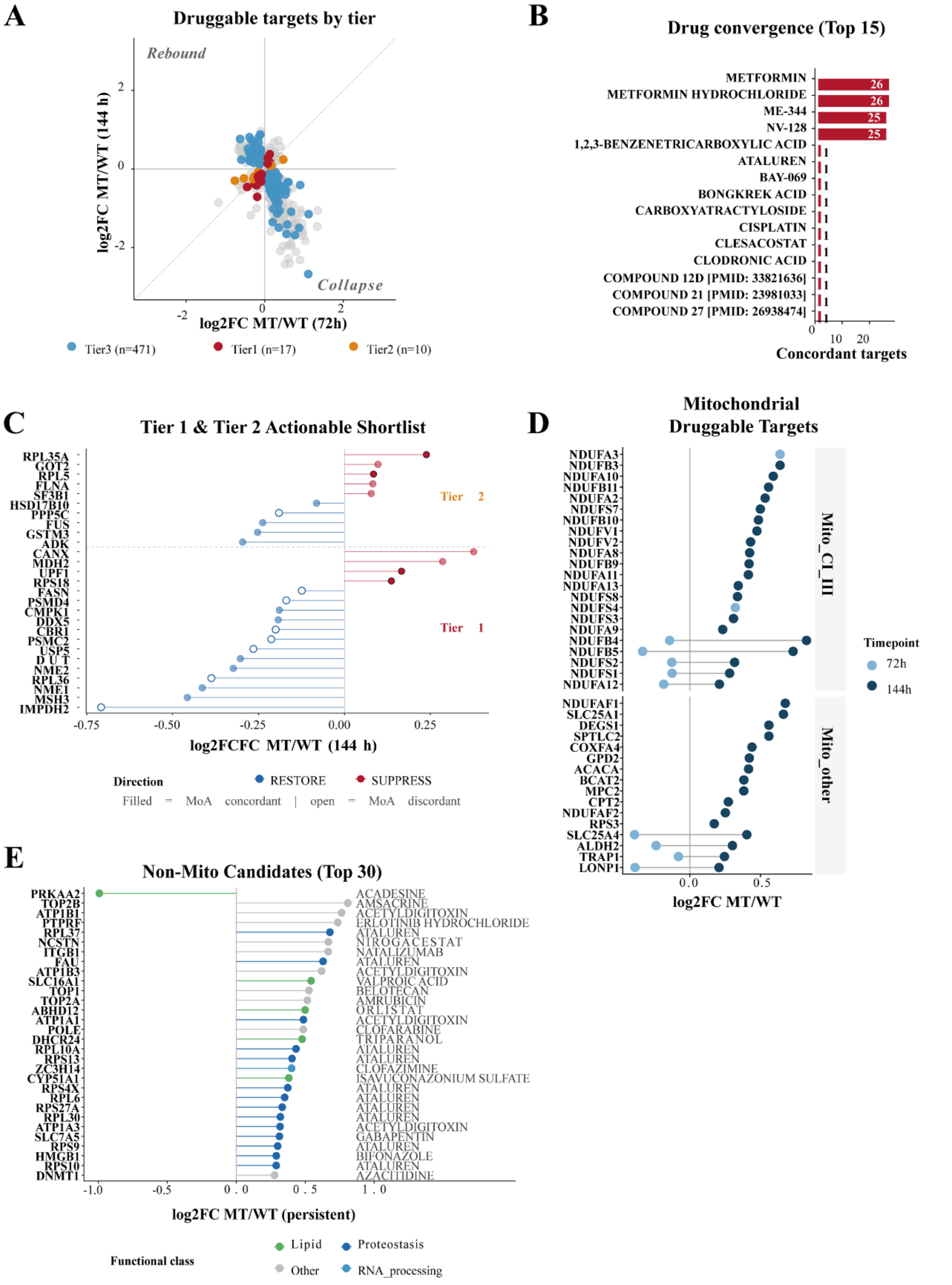


**Supplementary Figure 5. Extended direction-aware druggability screen: priority tiers, drug convergence and actionable candidates.** (A) Druggable targets across the biphasic landscape: every DEP plotted by $log_2FC$ (MT/WT) at 72 h versus 144 h and coloured by priority tier over the non-druggable background (grey, n = 1,213); Tier 1 (monotonic progressive trajectory, n = 17), Tier 2 (stable trajectory, n = 10) and Tier 3 (biphasic or single-time-point trajectory, n = 471). The number of Tier-3 targets that map to agents listed as approved in the source databases (361 of 471) is reported in Supplementary Table S6. (**B**) Drug convergence on mitochondrial targets (top 15 agents by number of concordant mitochondrial targets): metformin and metformin hydrochloride (26 targets each), ME-344 and NV-128 (25 each) far exceed all other agents, which map to a single target. (**C**) Tier 1 and Tier 2 actionable shortlist (27 targets) ordered by $\log_2$FC (MT/WT) at 144 h and coloured by the required correction (RESTORE, target depleted in mutant, blue; SUPPRESS, target elevated in mutant, red); filled points are mechanism-concordant and open points mechanism-discordant, with the Tier 1/Tier 2 boundary marked. (**D**) Mitochondrial druggable targets — temporal profile: $\log_2$FC (MT/WT) at 72 h (light) and 144 h (dark) for Mito_CI_III and Mito_other targets, including single-time-point proteins (Only72/Only144) that are not visible on the biphasic scatter. (**E**) Actionable non-mitochondrial candidates (top 30): concordant, approved agents complementing the mitochondrial arm, coloured by functional class (Lipid, Other, Proteostasis, RNA_processing) and labelled with the highest-phase drug ($\log_2$FC MT/WT, persistent).

A

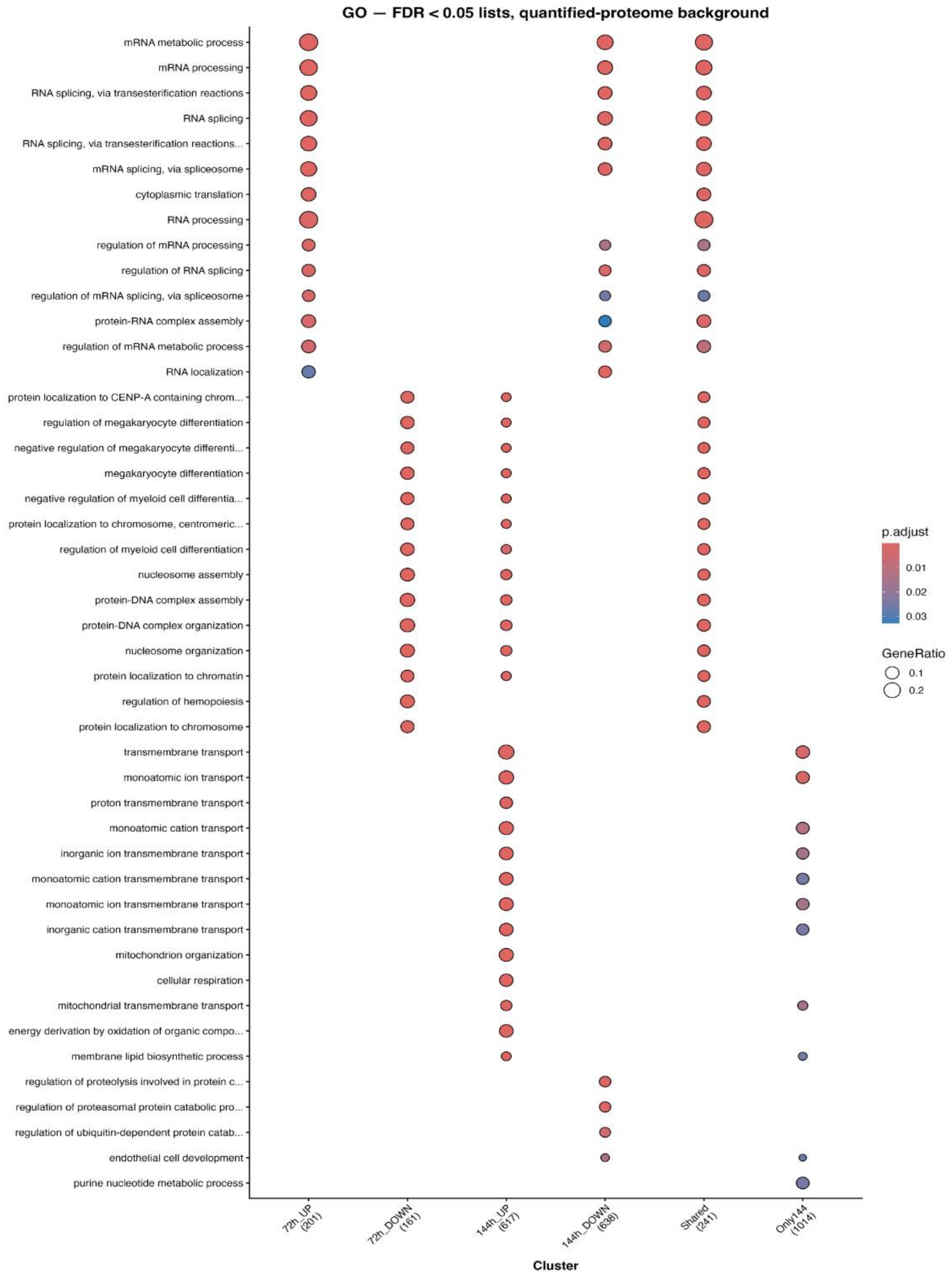
GO — FDR < 0.05 lists, quantified-proteome background
mRNA metabolic process
mRNA processing
RNA splicing, via transesterification reactions
RNA splicing
RNA splicing, via transesterification reactions...
mRNA splicing, via spliceosome
cytoplasmic translation
RNA processing
regulation of mRNA processing
regulation of RNA splicing
regulation of mRNA splicing, via spliceosome
protein-RNA complex assembly
regulation of mRNA metabolic process
RNA localization
protein localization to CENP-A containing chrom...
regulation of megakaryocyte differentiation
negative regulation of megakaryocyte differenti...
megakaryocyte differentiation
negative regulation of myeloid cell differentia...
protein localization to chromosome, centromeric...
regulation of myeloid cell differentiation
nucleosome assembly
protein-DNA complex assembly
protein-DNA complex organization
nucleosome organization
protein localization to chromatin
regulation of hemopoiesis
protein localization to chromosome
transmembrane transport
monoatomic ion transport
proton transmembrane transport
monoatomic cation transport
inorganic ion transmembrane transport
monoatomic cation transmembrane transport
monoatomic ion transmembrane transport
inorganic cation transmembrane transport
mitochondrion organization
cellular respiration
mitochondrial transmembrane transport
energy derivation by oxidation of organic compo...
membrane lipid biosynthetic process
regulation of proteolysis involved in protein c...
regulation of proteasomal protein catabolic pro...
regulation of ubiquitin-dependent protein catab...
endothelial cell development
purine nucleotide metabolic process
72h_UP (201)
72h_DOWN (161)
144h_UP (617)
144h_DOWN (638)
Shared (241)
Only144 (1014)
Cluster
p.adjust
0.01
0.02
0.03
GeneRatio
0.1
0.2

B

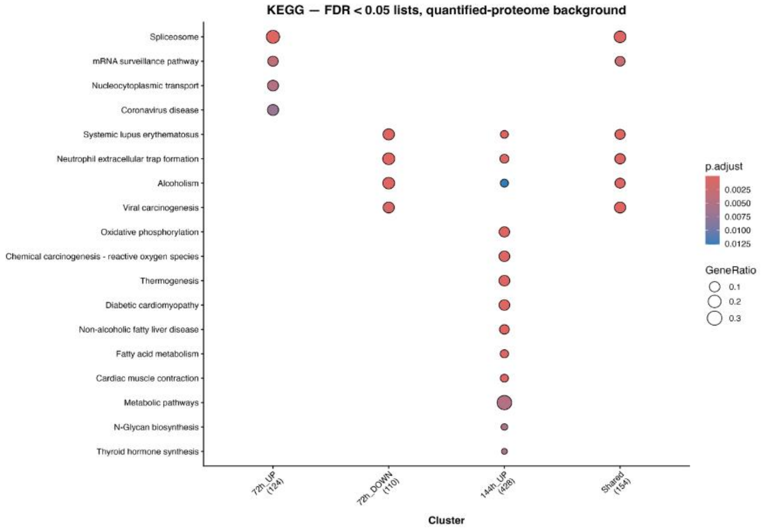
KEGG — FDR < 0.05 lists, quantified-proteome background
Spliceosome
mRNA surveillance pathway
Nucleocytoplasmic transport
Coronavirus disease
Systemic lupus erythematosus
Neutrophil extracellular trap formation
Alcoholism
Viral carcinogenesis
Oxidative phosphorylation
Chemical carcinogenesis - reactive oxygen species
Thermogenesis
Diabetic cardiomyopathy
Non-alcoholic fatty liver disease
Fatty acid metabolism
Cardiac muscle contraction
Metabolic pathways
N-Glycan biosynthesis
Thyroid hormone synthesis
72h_UP (124)
72h_DOWN (110)
144h_UP (428)
Shared (154)
Cluster
p.adjust
0.0025
0.0050
0.0075
0.0100
0.0125
GeneRatio
0.1
0.2
0.3

**Supplementary Figure 6. Sensitivity analysis of functional over-representation with FDR-filtered DEP lists and a quantified-proteome background. (A)** GO:BP and **(B)** KEGG over-representation (clusterProfiler compareCluster) for DEP sets defined at Benjamini–Hochberg-adjusted $p < 0.05$ (72h_UP, n = 209; 72h_DOWN, n = 161; 144h_UP, n = 629; 144h_DOWN, n = 655) and for the temporal classes (Only72, n = 123; Shared, n = 247; Only144, n = 1,037), using all proteins quantified at either time point as background. Only sets with at least one significant term are shown: Only72 had no significant GO:BP or KEGG term, and 144h_DOWN and Only144 had no significant KEGG term. Numbers under the x-axis give the genes of each set mapped to the annotation. The top-ranked terms per set (Benjamini–Hochberg-adjusted $p < 0.05$) are shown; dot size = gene ratio, colour = adjusted p-value; long GO:BP term names are truncated. Complete results are given in Supplementary Table S3 (REV_ENR sheets).
